# Oncofusion-driven *de novo* enhancer assembly promotes malignancy in Ewing sarcoma *via* aberrant expression of the stereociliary protein LOXHD1

**DOI:** 10.1101/2021.02.22.432287

**Authors:** Qu Deng, Ramakrishnan Natesan, Florencia Cidre-Aranaz, Shehbeel Arif, Ying Liu, Reyaz ur Rasool, Pei Wang, Zvi Cramer, Margaret Chou, Chandan Kumar-Sinha, Kristy Weber, T S Karin Eisinger-Mathason, Nicolas Grillet, Thomas Grünewald, Irfan A. Asangani

## Abstract

Ewing Sarcoma (EwS) is a highly aggressive tumor of bone and soft tissues that mostly affects children and adolescents. The pathognomonic oncofusion EWSR1-ETS (EWSR1-FLI1/EWSR1-ERG) transcription factors drive EwS by orchestrating an oncogenic transcription program through *de novo* enhancers. Pharmacological targeting of these oncofusions has been challenged by unstructured prion-like domains and common DNA binding domains in the EWSR1 and ETS protein, respectively. Alternatively, identification and characterization of mediators and downstream targets of EWSR1-FLI1 dependent or independent function could offer novel therapeutic options. By integrative analysis of thousands of transcriptome datasets representing pan-cancer cell lines, primary cancer, metastasis, and normal tissues, we have identified a 32 gene signature (ESS32 - Ewing Sarcoma Specific 32) that could stratify EwS from pan-cancer. Of the ESS32, LOXHD1 – that encodes a stereociliary protein, was the most exquisitely expressed gene in EwS. CRISPR-Cas9 mediated deletion or silencing of EWSR1-FLI1 bound upstream *de novo* enhancer elements in EwS cells led to the loss of LOXHD1 expression and altered the EWSR1-FLI1, MYC, and HIF1α pathway genes, resulting in decreased proliferation and invasion *in vitro* and *in vivo*. These observations implicate LOXHD1 as a novel biomarker and a major determinant of EwS metastasis and open up new avenues for developing LOXHD1-targeted drugs or cellular therapies for this deadly disease.

## Introduction

Ewing sarcoma (EwS) is the second most common malignant bone or soft-tissue cancer predominantly affecting children and young adults (1). Although the 5-year survival rate for primary EwS initially improved following the introduction of systemic chemotherapy in the neoadjuvant and adjuvant setting, several clinical studies indicate a plateau phase for these conventional therapies (2). Further, the prognosis for patients with high-risk recurrent disease is abysmal with <10% survival at 5 years; therefore, novel therapies are urgently needed to improve outcomes (1,3,4). EwS is driven by chromosomal translocations that generate pathognomonic fusions between the EWSR1 gene with variable members of the ETS family of transcription factors, most commonly FLI1 (85% of cases) (5,6). In the remaining 15–20% of EwS that are negative for EWSR1–FLI1 fusions, variant fusions between EWSR1 and other members of the ETS family occur, most commonly ERG (6).

EWSR1-FLI1 functions as a pioneer transcription factor by preferentially binding to genomic regions enriched for polymorphic GGAA microsatellites and induces chromatin reorganization resulting in the formation of opportunistic *de novo* enhancers and super-enhancers (7,8). Specifically, EWSR1-FLI1 binding to GGAA microsatellite repeats leads to the recruitment of the BRG1–BRM associated factor (BAF) chromatin-remodeling complex (9), BRD4 chromatin readers (10), lysine-specific demethylase (LSD1) (11) and RNA PolII (12), resulting in the establishment of *de novo* enhancers and activation of EwS transcriptional program (1,13). Although EWSR1-FLI1 would in principle constitute the most obvious and highly specific therapeutic target; this oncofusion protein represents a drug discovery challenge, because of its activity as targeting transcription factor and unstructured prion-like domains in the EWSR1 portion of the fusion (9,14). Given the paucity of druggable targets, the identification of novel Ewing-specific oncogene and mediators of EWSR1-FLI1 are urgently required.

Here, we use an integrative RNA-sequencing-based approach, coupled with ChIP-sequencing and tumor cell functional studies, to identify stereociliary protein LOXHD1 as a gene product specifically expressed in EwS. LOXHD1 meets the criteria for a potential oncogene, a diagnostic marker and exquisitely specific tumor antigen for potential adoptive cell-based therapy. Using various high throughput sequencing and functional studies, we have demonstrated that LOXHD1 is transcribed through EWSR1-FLI1 binding to an upstream *de novo* GGAA microsatellite with enhancer-like properties. Very little is known about the role of LOXHD1 in normal cell physiology or cancer due to its undetectable expression in a large majority of normal or cancer cells. To our knowledge, this is the first report studying the function of LOXHD1 outside the inner ear. Our studies implicate LOXHD1 as a major determinant of EwS metastasis through its ability to impact cytoskeletal reorganization, regulate EWSR1-FLI1, MYC transcription function, and hypoxic response through modulation of hypoxia-inducible factor 1α (HIF1α) stability. Together, our work provides strong evidence for LOXHD1 acting as an oncogene and even a potential cell-based immunotherapeutic target in EwS.

## Results

### Discovery of *LOXHD1* as an exquisitely specific EwS target gene

To identify highly specific EWSR1-FLI1 target genes with potential oncogenic function, we performed an integrative analysis of various ChIP-seq and transcriptomic datasets and mined for target genes uniquely expressed in EwS. The transcriptomic datasets included cancer cell lines (Cancer Cell Line Encyclopedia, CCLE, n = 980), normal tissues (Genotype-Tissue Expression, GTEx V6p, n = 11401), primary tumors (The Cancer Genome Atlas, TCGA, n = 9205), and pan-cancer metastatic tumor biopsies (MI-ONCOSEQ Program, MET500 cohort, n = 507) (15). A schematic of this pipeline is shown in **Fig. 1A**. Using cancer cell line RNA-seq, we first identified 516 genes that were commonly downregulated (>1 FPKM expression and ≥2-fold down) by EWSR1-FLI1 knockdown in three well-characterized EwS cell lines, SK-N-MC, A673, and CHLA-10 (8,10). We next used the CCLE dataset to filter out genes with expression >1 FPKM in any non-EwS cancer cell lines and narrowed our list to 89 highly specific EwS expressed genes (**Supplementary fig. S1A**). To qualify these genes as direct EWSR1-FLI1 transcriptional targets, we used ChIP-seq data for EWSR1-FLI1 enrichment in A673, SK-N-MC, and EWSR1-FLI1-overexpressing mesenchymal stem cells (MSCs), which are believed to be the cell-of-origin for EwS (**Supplementary fig. S1B**) (8). Genes that contain at least one EWSR1-FLI1 enrichment peak within ±100 kb of their TSS were selected as candidate direct targets. Our analysis identified 32 EwS-specific, EWSR1-FLI1 regulated genes, henceforth called ESS32 (**table S1**). Nearly all the genes in this set displayed a pronounced loss of expression upon knockdown of EWSR1-FLI1 in EwS cell lines and a gain of expression in MSCs ectopically overexpressing EWSR1-FLI1 (**Fig. 1B-left**). As an oncogenic pioneer transcription factor, EWSR1-FLI1 binds specifically to GGAA microsatellites repeat sequence and creates *de novo* enhancers from a closed chromatin conformation leading to transcriptional activation of multiple oncogenes (7–9). Interestingly, 75% of the EWSR1-FLI1-bound regions associated with ESS32 contained at least 5 consecutive GGAA microsatellite repeats (**Fig. 1B-right, Supplementary fig. S1C**), indicating strong EWSR1-FLI1 localization at these regions. To determine if these EWS-FLI-bound regions for ESS32 are *de novo* enhancers, we analyzed the presence of active enhancer mark, H3K27ac, in A673, SK-NM-C, primary EwS tissues, and MSCs (**Fig. 1C**). H3K27ac was significantly depleted around the ESS32 enhancers in both A673 and SK-N-MC upon EWSR1-FLI1 knockdown and was enriched in MSCs upon overexpression of EWSR1-FLI1; thus, demonstrating that the FLI1-bound enhancers are indeed formed *de novo*. In addition, the three primary EwS tissues showed an enrichment in H3K27ac levels, which are comparable to that in A673 and SK-N-MC cells. To illustrate the EwS-specific nature of ESS32 and its potential as a biomarker for EwS diagnosis, we analyzed the correlation of ESS32 expression among the samples in the MET500 dataset consisting of pan-cancer metastasis, and 11 EwS metastasis (**Fig. 1D**). The dense connectivity between 10 out of 11 EwS samples (Pearson correlation coefficient r>0.5), and the lack of such strong connectivity with other cancer types supports a high specificity of the ESS32 gene set to EwS. Furthermore, ESS32 gene signature stratified EwS from pan-cancer cohort in a Gene Set Enrichment Analysis (GSEA) (p-value <0.05, NES>1.5) demonstrating its power and specificity toward EwS, which could be used as diagnostic marker (**Fig. 1E**). The ESS32 gene set comprised of some known EwS genes such as KLF15, NKX2-2, STEAP2, etc. as well as many new targets that have not yet been associated with EwS (**table S1**). However, it still presented a weak degree of association with other cancers (prostate cancer (PRAD), Cholangiocarcinoma (CHOL), and ovarian cancer (OV). Next, we filtered out genes contributing to this non-specificity using an additional filter based on the TCGA and MET500 datasets, wherein we discarded any genes with >1 FPKM expression in more than one cancer type (**Supplementary fig. S1D** and **S1E**). This narrowed the ESS32 list to three EwS-specific genes, namely *RBM11, LIPI*, and *LOXHD1*, of which *RBM11* displayed high expression across multiple tissue types in the GTEx dataset, whereas LIPI and LOXHD1 showed marginal expression only in thyroid and testis, respectively (**Supplementary fig. S1F**). Both *LOXHD1* and *LIPI* showed exquisitely restricted expression in the metastatic EwS tissues and in none of the other pan-metastatic tissues (**Fig. 1F** and **Supplementary fig. S1G**). Remarkably, the specificity of *LOXHD1* and *LIPI* surpassed the commonly used EwS diagnostic marker *CD99, NKX2-2, PAX-7, BCL11B*, and *GLG1* (16–19)(**Fig. 1G** and **Supplementary fig. S1G**). *LIPI* has been reported to be upregulated in EwS where it is known to regulate lysophosphatidic acid mediated signaling (20,21). However, little is known about LOXHD1 expression and function in normal physiology and cancer. To further define the spectrum of LOXHD1 expression in EwS, we queried the publicly available Affymetrix array data comprising normal, primary, metastasis, and recurrent EwS samples (22); and found its expression to be significantly upregulated in primary tissue, and associated with disease progression (**Supplementary fig. S1H**). To further validate these findings, we profiled 12 primary EwS patient samples for *LOXHD1* using qRT-PCR, of which 10 EwS samples displayed over a hundred-fold higher *LOXHD1* expression compared to the 2 non-EwS samples (**Supplementary fig. S1I**). These data strongly suggesting *LOXHD1* as a highly specific EwS gene, led us to investigate the role of LOXHD1 as an oncogene mediator driving EwS.

**Figure 1:**
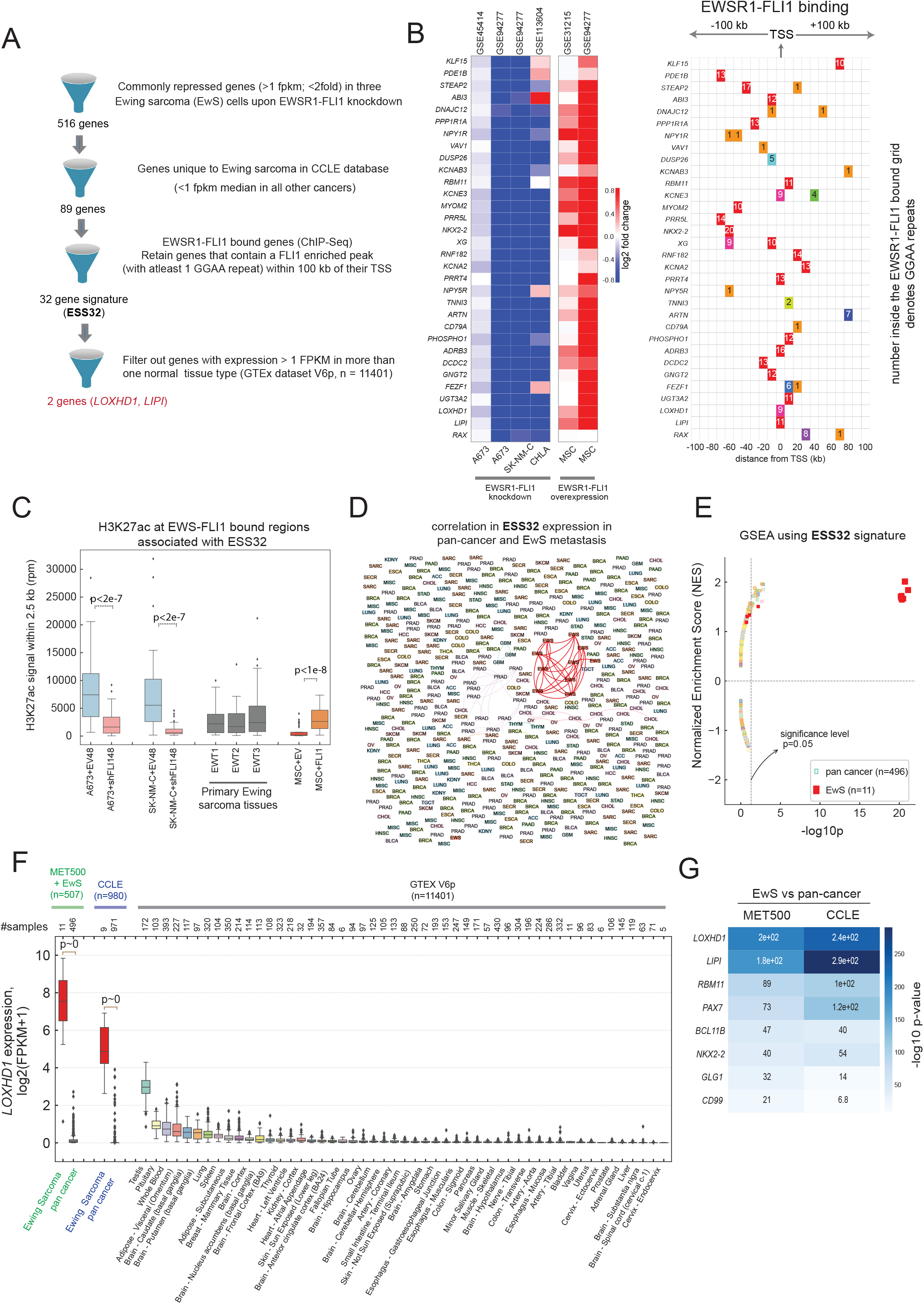
Integrative analysis leading to the identification of ESS32 gene signature, and stereociliary protein LOXHD1 as EwS specific gene. **(A)** Flowchart depicting the various stages of analysis in our computational pipeline to identify direct EWS-FLI targets in EwS. **(B)** *(left)* log2 fold change in expression (microarray and RNA-seq), for EwS specific 32 genes signature (ESS32) showing their marked downregulation in EWSR1-FLI1-knockdown EwS cell lines (n=4), and marked upregulation in EWSR1-FLI1-overexpressing Mesenchymal Stem Cells (MSC, n=2), *(right)* table showing the positions of EWSR1-FLI1-bound (ChIP-seq) regulatory regions within ±100 kb of transcription start-site (TSS); the numbers denote the number of polymorphic GGAA-microsatellite-repeats contained within each regulatory element. **(C)** Increased transcription activation mark H3K27ac on ESS-32 regulatory region. Total H3K27ac tag density (rpm) within ±2.5 kb of the EWSR1-FLI1 bound regulatory regions for ESS-32 is shown for two EwS cell lines upon EWSR1-FLI1 knockdown, three EwS primary tumor tissue and MSCs overexpressing EWSR1-FLI1. **(D)** ESS-32 expression identifies EwS among hundreds of pan-cancer metastatic disease. Network plot for the correlation in the expression of ESS-32 genes in RNA-seq data of MET500 pan-cancer metastatic tumor biopsies (n= 500) and metastatic EwS (n= 11). Connectivity is displayed only for samples with Pearson-correlation-coefficient *r* ≥ 0.5 and the thickness of the connections is proportional to *r*. Here EWS, PRAD, CHOL, OV, SARC denote EwS, prostate cancer, cholangiocarcinoma, ovarian cancer and sarcoma subtypes, respectively. Complete abbreviation of sample names can be found in SI. **(E)** GSEA analysis of ESS-32 geneset in MET500 and Ewing sarcoma RNA-seq data, showing its significant enrichment in >70% of EwS metastatic samples. **(F)** LOXHD1 is predominantly expressed only in EwS tumors and tumor-derived cell lines. *LOXHD1* expression in MET500 and EwS metastatic biopsies (n=507), Cancer Cell line encyclopedia (CCLE, n=980) and Genotype tissue expression (GTEX, n=11401) transcriptomic datasets. **(G)** Heatmap shows the log-transformed p-values computed for EwS vs pan-cancer samples in MET500+EwS (n=507) and CCLE (n=980) datasets, for the mentioned genes, see fig. S1G. The p-values were computed using an independent, two-sample t-test.

### Re-annotation and mapping of LOXHD1 transcript and protein in EwS cells

*LOXHD1* codes for an evolutionarily conserved protein predicted to contain 15 PLAT (polycystin-1, lipoxygenase,alpha-toxin) domains. The biological function of PLAT domains is not well established, but it is speculated that they target proteins to the plasma membrane (23,24). Very little is known about the role of LOXHD1 in normal cell physiology or in cancer, due to its undetectable expression in a large majority of normal or cancer cells. Therefore, we first sought to identify the gene and protein structure of the *LOXHD1* in the EwS cells. The Ensembl GRCh37 annotation of *LOXHD1* gene (ENSG000000167210, chr18: 44056935-44236996) shows 40 exons and multiple splice isoforms. The 6848 bp long major isoform ENST00000536736 encodes the canonical 2211 amino acids (aa) protein (UniprotKB ID: F5GZB4) which contains 15 PLAT/LH2 and one coiled-coil domain. However, our analysis of RNA-seq data from 12 EwS cell lines showed nearly zero transcriptional output for the first seven exons (**Fig. 2A**), suggesting that exons 1 through 7 in the current annotations may not be part of the *LOXHD1* gene structure in the EwS cells. To test this hypothesis, we integrated ChIP-seq data H3K4me3 (marker associated with TSS) and H3K27ac (active transcription mark), with RNA-seq from 12 EwS cell lines to map the EwS specific *LOXHD1* transcript structure. A common TSS for *LOXHD1* was found in all of the twelve EwS cell lines located at the 8th exon in the current Ref. Seq. annotation (**Fig. 2A**). ChIP-seq tracks revealed that the active enhancer/promoter and TSS region for *LOXHD1* locates slightly upstream of exon 8, as evidenced by the pronounced enrichment of both H3K27ac and H3K4me3 mark, respectively in the region around exon 8 (**Fig. 2A**). Additionally, the ENCODE DNase hypersensitivity data for two EwS cell lines A673 and SK-N-MC also showed a DNAse 1 hypersensitive region near our newly annotated H3K4me3 bound TSS, and H3K27ac bound regulatory elements, further validating our observation that the *LOXHD1* transcription in EwS proceeds through an alternate start site with a potential upstream regulatory element. (**Fig. 2A**). The remapped transcript structure contains 33 exons and codes for 1891 aa protein containing 13 PLAT and 1 coiled-coil domains (**Fig. 2B**). We predicted the coiled-coil structure for amino-acids 596 through 658 in the remapped protein using the COILS server (25) (**Fig. 2B and Supplementary fig. S2A**). Additionally, using the NLS (nuclear localization signal) mapper (26), we identified a cryptic NLS (aa 616-629) within the coiled-coil region (**Supplementary fig. S2B**). Immunoblot analysis in a panel of EwS (n=3) and non-EwS (n=3) cell lines using LOXHD1 specific antibody (23) displayed a specific band between 200-220 kDa in EWSR1-FLI1 expressing cells (**Fig. 2C**), which is consistent with the predicted molecular weight of a 1891 aa (216.4 kDa) LOXHD1 protein. The presence of coiled-coil and NLS domain suggested nuclear localization of LOXHD1 in addition to its cytosolic functions. To test whether LOXHD1 localizes to the nucleus, we cloned a portion of mouse *Loxhd1* exon19 which is homologous to human *LOXHD1* with over 92% sequence similarity and encompasses both the NLS and coiled-coil domains (**Supplementary fig. S2C**). Overexpression of the HA-tagged LOXHD1 exon19 in 293T cells showed strong immunofluorescence staining in the cell nucleus (**Supplementary fig. S2C**). We further confirmed this feature through immunofluorescence staining of the endogenous LOXHD1 in the EwS cell lines. In SK-N-MC and RDES cells, LOXHD1 staining was observed on both the plasma membrane and in the nucleus, whereas prostate cancer cell line LNCaP used as a negative control displayed no specific staining (**Fig. 2D**). Together, these data demonstrate an alternative transcription start site for *LOXHD1* in EwS cells and provide evidence for its protein expression in both cytoplasmic and nuclear compartments.

**Figure 2:**
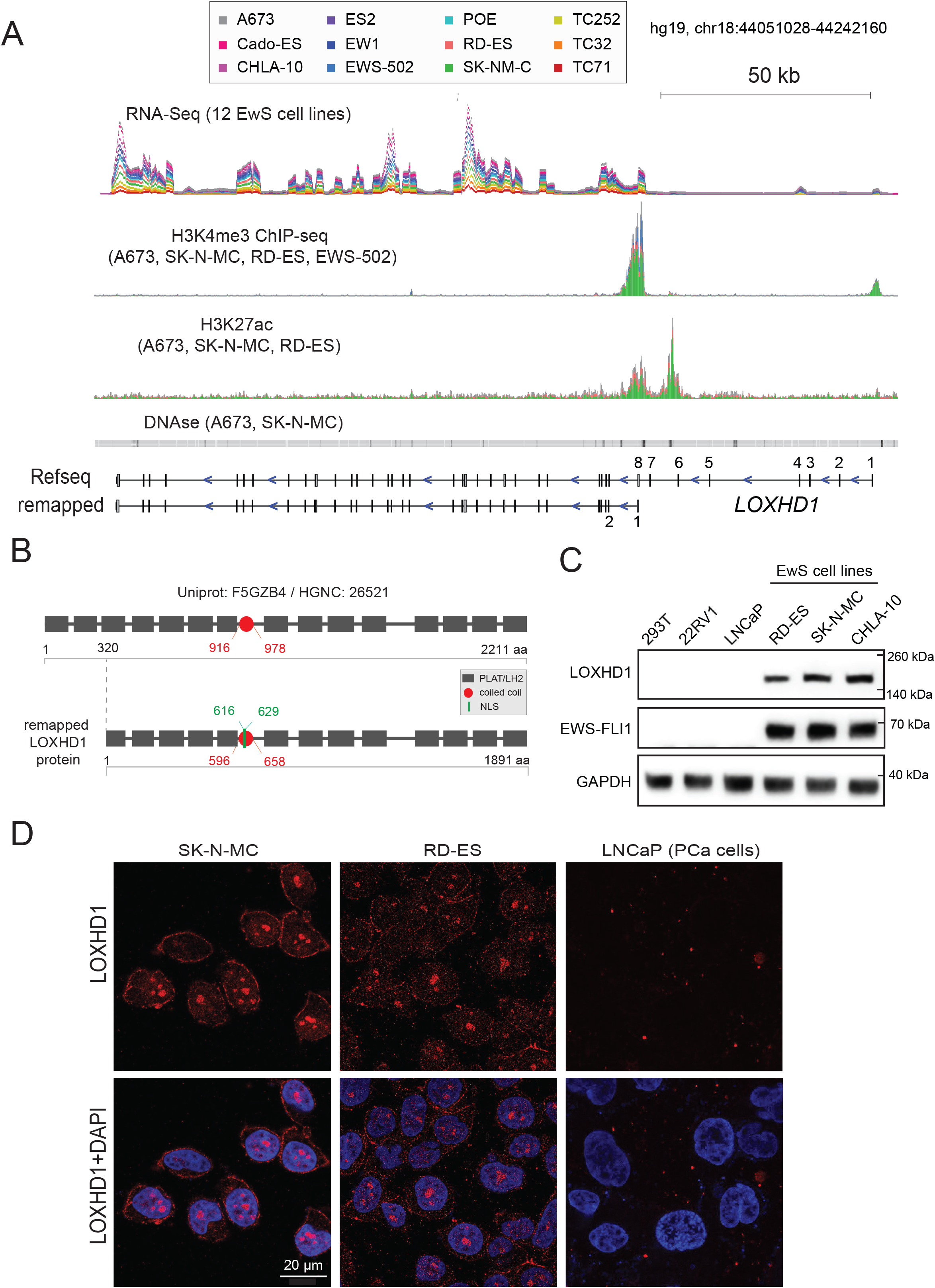
Alternative TSS and identification of NLS in LOXHD1: **(A)** IGV plot of RNA-seq in multiple EwS cell lines, along with ChIP-seq track for transcription activation mark H3K4me3 and H3K27 showing alternative Transcription Start Site (TSS) for *LOXHD1* in EwS cells. ENCODE DNase 1 hypersensitivity (HS) data for SK-N-MC and A673, showing HS site near the TSS. **(B)** Protein domain structure of LOXHD1. *Top*, based on ref seq. *Bottom*, based on transcript excluding first seven exon sequences as because of alternative TSS. The protein is composed of thirteen PLAT domain. Newly identified NLS (nuclear localization signal) and the coiled-coil domain is indicated with aa position. **(C)** Detection of stereociliary LOXHD1 protein in EwS cells. Immunoblot analysis for LOXHD1 and EWSR1-FLI1 levels in three EwS cells and two prostate cancer cells (LNCaP, 22RV1) and HEK293T cells. GAPDH used as a loading control. **(D)** Immunofluorescence imaging showing LOXHD1 (red) expression on the plasma membrane and in the nucleus in RD-ES and SK-N-MC cells. DAPI (blue) used to stain the nucleus. LNCaP cells were used as a negative control for LOXHD1 expression.

### EWSR1-FLI1 binding to the GGAA microsatellite creates *de novo* enhancer upstream of LOXHD1 and regulates its expression

As a pioneer transcription factor, EWSR1-FLI1 binds specifically to GGAA microsatellites repeat sequence and creates *de novo* enhancers from a closed chromatin conformation leading to transcriptional activation of multiple oncogenes (7–9). In **Fig.1A-C**, we showed that the entire **ESS32** gene set presents EWSR1-FLI1 binding to distal *de novo* enhancer sites, which contains at least 5 GGAA repeats within 100 kb of their TSS. We identified the regulatory region for *LOXHD1* located roughly 6.7 kb upstream of the newly annotated TSS and contained 9 consecutive GGAA repeats. To demonstrate the presence and *de novo* nature of enhancer regulating *LOXHD1* expression, we analyzed ChIP-seq data (8) for MSCs in control and EWSR1-FLI1 overexpression conditions (**Fig. 3A**). Ectopic expression of EWSR1-FLI1 displayed *de novo* enhancer formation characterized by active chromatin H3K27ac mark and a strong EWSR1-FLI1 peak ∼6.7 kb upstream from the *LOXHD1* TSS; and transcription of *LOXHD1* (**Fig. 3A and Supplementary fig. S3A**). As expected, the same site was occupied by endogenous EWSR1-FLI1 with enriched H3K27ac mark in SK-N-MC cells (**Fig. 3A**). Similar to this, a *de novo* enhancer spaced between *LIPI* and *RBM11* was observed – which might regulate their expression (**Supplementary fig. S3C**). These observations indicate that LOXHD1, which is transcriptionally silent in a vast majority of normal and pan cancer cells, is induced exclusively in EwS by the pathognomonic oncofusion EWSR1-FLI1. Consistent with these findings, infection of U2OS osteosarcoma cells with EWSR1-FLI1 led to *LOXHD1* expression (**Fig. 3B**). Likewise, in an orthogonal approach, EWSR1-FLI1 knockdown in SK-N-MC and A673 cells resulted in the disassembly of the *LOXHD1* enhancer with a complete loss of H3K27ac mark and a resulting loss in the expression of *LOXHD1* (**Fig. 3C**). To further validate the role of EWS-ETS in regulating *LOXHD1* expression, we knocked down EWSR1-FLI1 or EWS-ERG by shRNA in a panel of EwS cells and found 2-10 fold decrease in *LOXHD1* expression by qRT-PCR analysis (**Fig. 3D and Supplementary fig. S3B**). Additionally, inhibiting BET bromodomain, that we previously demonstrated to be important for EWS-ETS mediated transcription (10)-with JQ1 in EwS cells resulted in the loss of *LOXHD1* expression (**Fig. 3E**), further indicating the role of EWS-ETS and its associated transcriptional complex in LOXHD1 transcription. Together, we established that *LOXHD1* is regulated by a distal EWSR1-FLI1 bound enhancer region located 6.7 kb upstream of its TSS. These observations demonstrate transcriptional regulation of *LOXHD1* through a distal *de novo* enhancer assembled by the pathognomonic EWS-ETS transcription factor in EwS, and also suggests that LOXHD1 is not expressed in the vast majority of normal and cancer cells.

**Figure 3:**
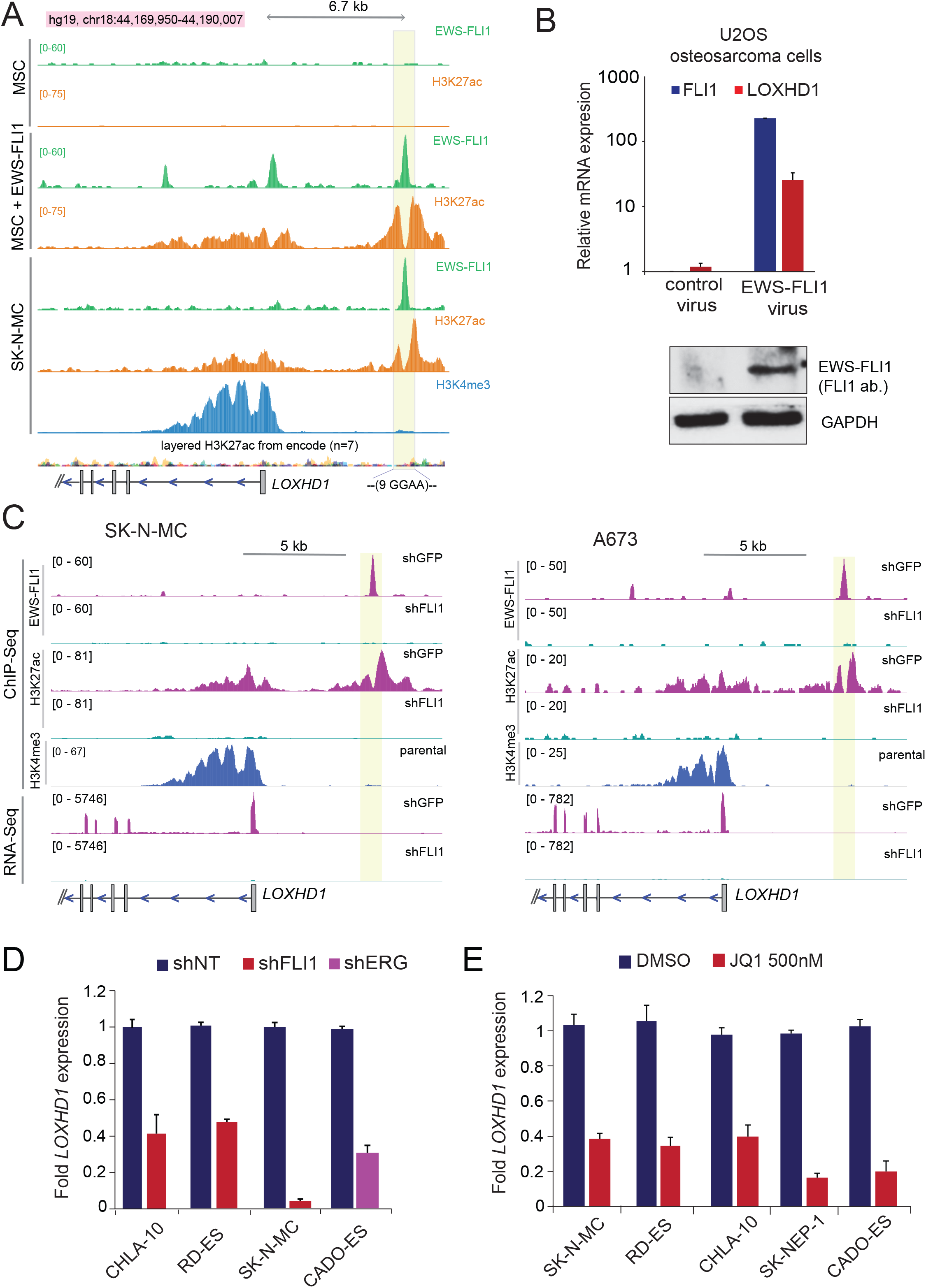
EWSR1-FLI1 binding to the polymorphic GGAA microsatellite creates a *de novo* enhancer upstream of *LOXHD1* and regulates its expression. **(A)** EWSR1-FLI1 binding upstream of *LOXHD1* creates a *de novo* enhancer. Genome browser view of ChIP-seq tracks showing the EWSR1-FLI1 binding to *LOXHD1* upstream region containing GGAA repeats, generating a *de novo* enhancer marked by H3K27ac mark in EWSR1-FLI1 overexpressing MSC, and wild type SK-N-MC cells (GEO accession code # GSE94278). **(B)** Ectopic expression of EWSR1-FLI1 leads to transcriptional activation of LOXHD1 in non-Ewing cancer cells. *top*, qRT-PCR showing upregulation of LOXHD1 in U2OS osteosarcoma cells upon EWSR1-FLI1 expression. *bottom*, Immunoblot for EWSR1-FLI1, and GAPDH (loading control) in the indicated samples. **(C)** Knockdown of EWSR1-FLI1 collapses the *de novo* enhancer leading to silencing of *LOXHD1* expression. Genome browser view of integrated ChIP-seq and RNA-seq tracks showing loss of EWSR1-FLI1 enrichment to the GGAA microsatellite with a concomitant loss of H3K27ac mark and transcriptional silencing of LOXHD1, respectively upon shFLI1 mediated EWSR1-FLI1 knockdown in SK-N-MC (*left*) and A673 cells (*right*). ChIP-seq track for H3K4me3 denotes the TSS (GEO accession code # GSE94278). **(D** and **E)** EWS-ETS fusion knockdown or inhibition of its co-activator BRD4 downregulates *LOXHD1* expression. qRT-PCR showing downregulation of *LOXHD1* transcript upon shRNA mediated EWSR1-FLI1/EWSR1-ERG knockdown or treatment with JQ1 at 500nM for 24h in a panel of EwS cell lines.

### Genomic deletion or epigenetic silencing of the EWSR1-FLI1 bound *de novo* enhancer represses *LOXHD1* transcription

We next investigated the functional association between *LOXHD1* expression and its EWSR1-FLI1 bound upstream enhancer by deleting the GGAA microsatellite repeats through CRISPR-Cas9 genome editing. Infection with sgRNA Cas9 lentivirus targeting regions on either side of the GGAA repeat led to the deletion of approximately 172bp DNA (**Fig. 4A** and **Supplementary fig. S4A**) and a concomitant decrease in the transcription of *LOXHD1* in a pool population of SK-N-MC and RD-ES cells (**Fig. 4B**). Additionally, we observed >90% reduction in *LOXHD1* mRNA levels in independent single clones containing the enhancer deletion (eKO1 and eKO2) (**Fig. 4B right** and **Supplementary fig. S4B-C**). In an orthogonal set of experiments, we silenced the activity of the *LOXHD1* enhancer using the CRISPR dCas9-KRAB system (CRISPRi) with two independent sgRNAs (eKD1 and eKD2) targeting the adjacent region of the EWSR1-FLI1 bound GGAA repeat (**Fig. 4C**). CRISPRi induces focal chromatin-repressive states by KRAB mediated H3K9me3 deposition at target sites (27,28). We first assessed and found the accumulation of H3K9me3 chromatin mark within the targeted microsatellite region in SK-N-MC and RD-ES cells transduced with CRISPR dCas9-KRAB constructs (**Fig. 4D**). Consistent with histone deacetylase activity of the KRAB domain, ChIP-seq analysis of H3K27ac mark demonstrated the loss of signal from the targeted microsatellite and adjacent region (**Fig. 4E**). Additionally, reduction in H3K4me3 and H3K27ac mark from the *LOXHD1* TSS was observed in CRISPRi cells compared to controls, suggesting that the chromatin state changes on the distal enhancer could affect the active transcription mark likely due the loss of enhancer-promoter contact (**Fig. 4E**). As expected, compared to control cells, repression of the enhancer by two independent gRNA led to significant loss of *LOXHD1* transcription (**Fig. 4F**). These results were further confirmed by immunoblotting and immunofluorescence analysis that demonstrated the reduction of LOXHD1 protein levels in the eKD polyclonal pools and eKO single cell clones compared to controls (**Fig. 4G** and **4H**). Altogether, our findings from genetic deletion and epigenetic silencing approaches provide substantial evidence that *LOXHD1* is a direct target of EWSR1-FLI1, and its expression is regulated by a distal EWSR1-FLI1 bound GGAA-rich *de novo* enhancer region in EwS.

**Figure 4:**
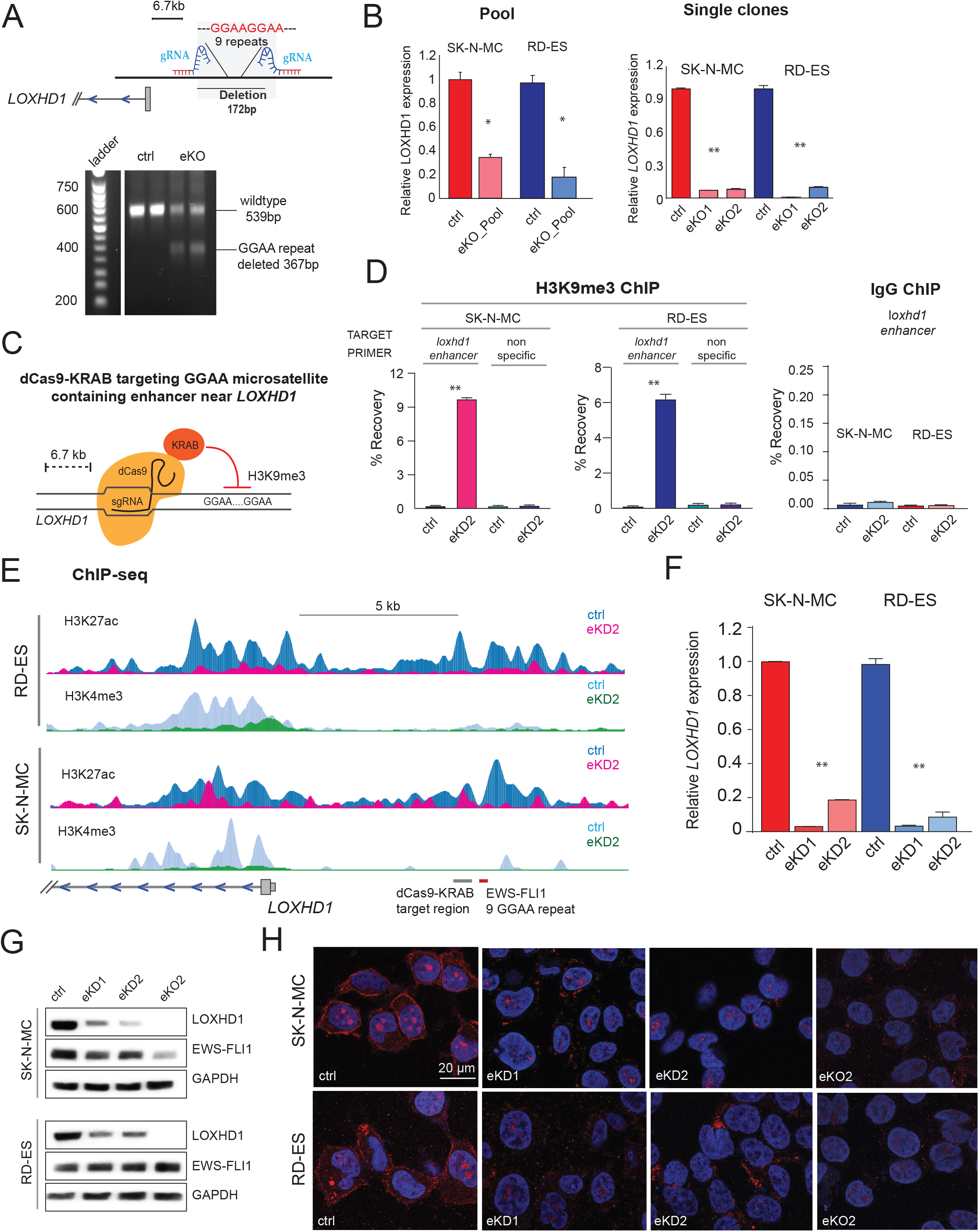
Knockout or dCas9-KRAB mediated silencing of EWSR1-FLI1 bound *de novo* enhancer quashes LOXHD1 transcription: **(A)** *above*, Schematic showing the CRISPR sgRNA specifically flanking the GGAA microsatellite upstream of *LOXHD1. Below*, DNA ethidium bromide stained gel image showing the deletion of GGAA repeats in SK-N-MC and RD-ES cells transduced with enhancer targeting sgRNA lentivirus or Cas9 control lentivirus. **(B)** qRT-PCR showing *LOXHD1* expression in enhancer knockout pools *(left)* and two independent isogenic single-cell clones *(right)* compared to their respective controls. **(C)** Schematic showing the CRISPR-dCas9-KRAB (CRISPRi) mediated epigenetic silencing of GGAA microsatellite containing *LOXHD1* enhancer. Two independent small guide RNAs (sgRNA1 and sgRNA2) were designed adjacent to the GGAA microsatellite. **(D)** ChIP-qPCR analysis showing accumulation of KRAB catalyzed H3K9me3 mark at the *LOXHD1* upstream GGAA microsatellite region (m.s. region primers) in SK-N-MC and RD-ES cells expressing dCas9-KRAB and sgRNA2 (eKD2 – enhancer KnockDown 2). A pair of non-specific (n.s.) region primers were used as negative control and IgG served as ChIP negative control. **(E)** Depletion of active transcription marks upon *de novo* enhancer knockdown. ChIP-seq tracks of H3K27ac and H3K4me3 signals at the *LOXHD1* loci in RD-ES and SK-N-MC cells expressing dCas9-KRAB and sgRNA2 as in E. gRNA target site and EWSR1-FLI1 target microsatellite region is indicated with thick gray and red line, respectively **(F)** qRT-PCR showing the loss of *LOXHD1* expression in enhancer knockdown cells. **(G)** Immunoblots showing loss of LOXHD1 protein in enhancer knockdown (dCas9-KRAB) or enhancer knockout (CRIPSR-cas9) cells. EWSR1-FLI1 and GADPH was used as control. **(H)** Immunofluorescent staining of LOXHD1 (red). The nucleus was visualized by DAPI (blue). *p < 0.05, **p < 0.001 by two-tailed Student’s t test.

### LOXHD1 silencing impairs major oncogenic transcription factor response and cytoskeletal organization

Except for a study showing that a missense mutation in mouse *LOXHD1* affects the function of the sensory cells involved in hearing (23), not much is known about its role in normal or cancer cell physiology. Towards that end, we first attempted to understand the consequence of LOXHD1 loss of expression in the EwS cells by performing RNA-seq experiment in parental versus enhancer knockdown (eKD) SK-N-MC and RD-ES cells. The enhancer KD cells displayed significant change in the transcriptome with hundreds of genes up and downregulated as a result of LOXHD1 silencing (**Supplementary fig. S5A**). GSEA following RNA-seq suggested that a majority of the 512 EWSR1-FLI1-target genes (**Fig.1 A**) and the **ESS32** gene set were both negatively enriched upon LOXHD1 silencing (**Fig. 5A *top*** and **Supplementary fig. S5B**). GSEA analysis further identified negative enrichment of oncogenic hallmark MYC signature in the LOXHD1 silenced SK-N-MC and RD-ES cells (**Fig. 5A *bottom***). Together it suggested the LOXHD1 silencing may have a negative effect on the tumorigenic potential of EwS cells. Additionally, Gene Ontology (GO) pathway enrichment analysis showed cytoskeleton organization and actin family protein among the top deregulated biological pathways (**Fig. 5B**). Plasma membrane-associated proteins regulate cytoskeletal assembly through their ability to regulate components of the actin and microtubule filament network (29). Reorganization of cytoskeleton affects cell signaling, polarity, motility, cell-cell, and cell-ECM (extra cellular matrix) interactions, and, more importantly, alters the metastatic potential of cancer cells (30,31). Based on the above observations and given the fact that LOXHD1 primarily was found to be associated with the plasma membrane (23), we hypothesized that LOXHD1 regulates EwS cell cytoskeleton and promotes tumorigenesis. Immunofluorescent staining of F-actin in RD-ES and SK-N-MC cells displayed a well-organized cytoskeletal structure underneath the plasma membrane with elongated nuclear morphologies representative of spread, adherent cells (**Fig. 5C**). However, LOXHD1 silenced eKD1 and eKD2 cells displayed diffuse, highly irregular cytoskeletal patterns with circular nuclear morphologies representative of non-adherent cells. The cell surface area, which is directly related to the degree of cellular adhesion to its substrate, for eKD1 and eKD2 cells, was substantially smaller compared to its controls (**Fig. 5D**). The data indicated that the silencing of LOXHD1 alters the cell-to-cell and cell to matrix interactions. We then hypothesized that cell growth at single-cell density which requires optimum cell-to-cell contact could be compromised in the eKD cells, and tested it by sphere formation on 3D matrigel, and 2D colony formation assays. As expected, LOXHD1 silenced cells form substantially less and small spheres and colonies than parental controls (**Fig. 5E** and **Supplementary fig. S5C**). We further tested the ability of eKD cells to form aggregates by suspending them in 24-well ultralow attachment plates. While the parental cells exhibited stabilized, large aggregates containing hundreds of cells, the eKD cells did not display similar aggregates even after 16 hours post-plating, suggesting a reduced cell-cell contact potential as a result of altered cytoskeleton in the LOXHD1 silenced cells (**Fig. 5F**). Since the cytoskeletal organization can significantly alter the migratory potential of cancer cells, we performed wound healing and Boyden chamber invasion assays. We found significantly reduced migration and invasion of LOXHD1 silenced cells compared to the respective parental controls (**Fig. 5G** and **Supplementary fig. S5D-5E**). Notably, we did not observe any change in the proliferation rates of LOXHD1 silenced cells in 2-D cell culture in this study (data not shown). Together, these data demonstrate that LOXHD1 silencing in EwS cells impairs major oncogenic transcription factor pathways including EWS-ETS, MYC and cytoskeletal organization, resulting in reduced anchorage independent growth and metastatic potentials *in vitro*.

**Figure 5:**
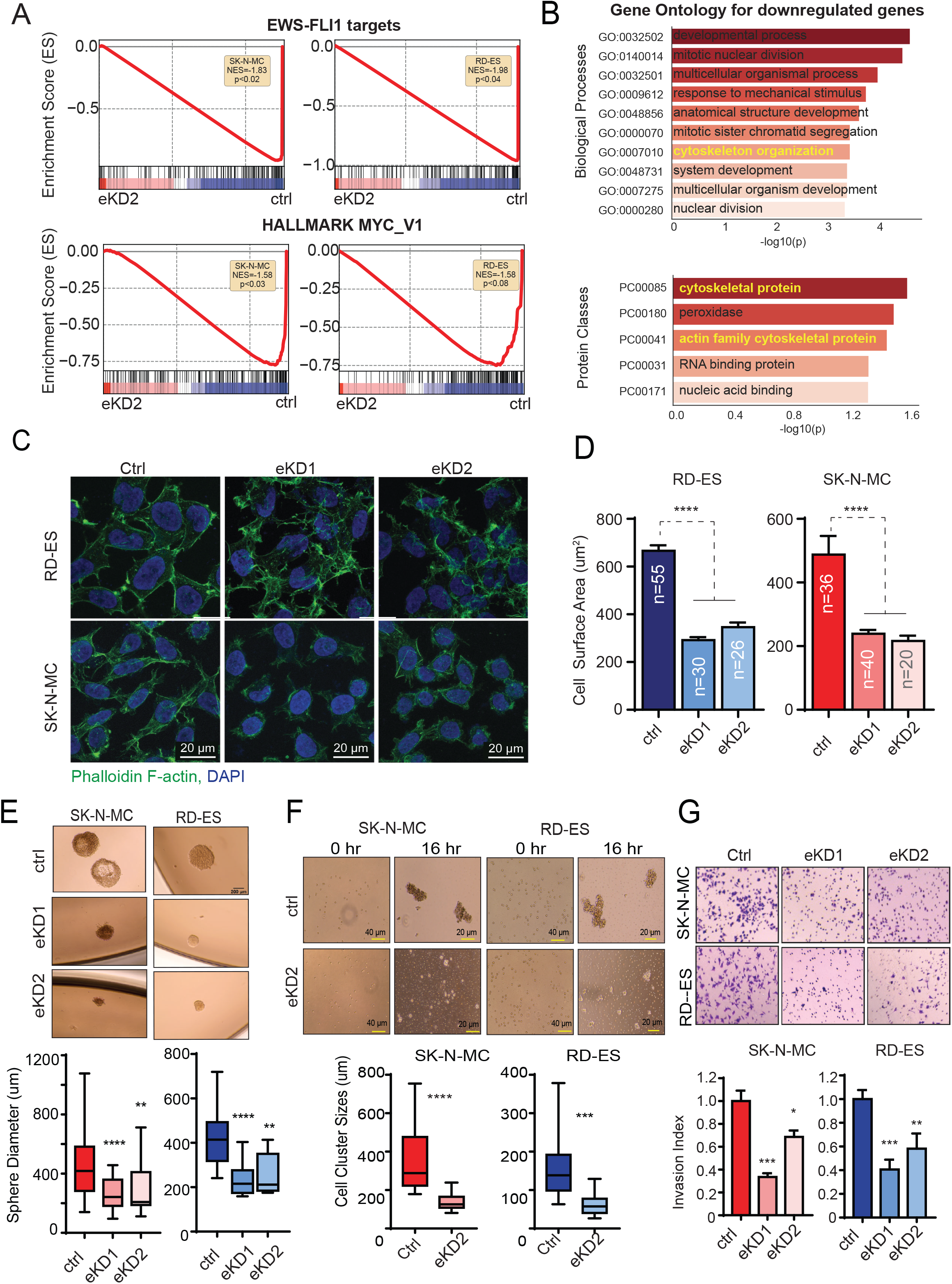
LOXHD1 loss impairs major oncogenic transcription factor response and cytoskeletal organization. **(A)** RNA-seq followed by GSEA showing negative enrichment of EWSR1-FLI1 and MYC gene signatures in the *LOXHD1* enhancer knockdown (eKD2) EwS cells. **(B)** Gene Ontology (GO) terms of biological process and protein classes for common downregulated genes in *LOXHD1* eKD2 cells. **(C-D)** Cytoskeletal disorganization in LOXHD1 silenced cells. **(C)** Representative images of Phalloidin F-actin and DAPI staining in RD-ES and SK-N-MC cells with and without LOXHD1 silencing. **(D)** Quantification of cell surface areas in the immunofluorescence staining images with imageJ. **(E)** LOXHD1 silenced cells display reduced anchorage-independent growth. *Top* Representative images of a sphere formation assay performed with indicated cells embedded in 50% of Matrigel in a 24 well plate. Quantification is shown in *bottom* panel. **(F)** LOXHD1 silenced cells display reduced cell aggregation property. *Top* Representative images of a cell aggregation assay performed by seeding single-cell suspension on poly-HEMA coated ultralow attachment plates. Images were taken at 0hr and 16hr. Quantification is shown in *bottom* panel. **(G)** LOXHD1 silenced cells display reduced Matrigel invasion. *Top* Representative images of invaded cells 48h post plating are shown for control and *LOXHD1* enhancer knockdown cells. *bottom* Quantification. **** p< 0.0001, ***p<0.001, **p<0.01, *p<0.05 by two-tailed Student’s t-test.

### LOXHD1 knockdown attenuates hypoxia response in EwS cells by destabilizing HIFtα

Intratumoral hypoxia is a common feature of solid malignancies including sarcomas. Hypoxia-inducible factor (HIF) proteins, mainly HIF1α and HIF2α, are transcription factors essential for cellular adaptation to hypoxic stresses (32). Overexpression of HIF1α has been shown to enhance the metastatic potential of sarcomas, including EwS, and other solid cancers (33–35). Since LOXHD1 protein was found in the nuclear compartment of the EwS cells, we wondered whether it has a role in HIF1α transcriptional output. Remarkably, the invasive potential of LOXHD1 proficient EwS cell was amplified when the Boyden chamber assays were conducted under hypoxic conditions (**Fig. 6A**, and **Supplementary fig. S6A**). The SK-N-MC cells displayed 2-fold higher invasion in hypoxic culture than normoxia (**Fig. 6A**), confirming earlier reports that hypoxia promotes sarcoma invasion and metastasis (33–35). In contrary, LOXHD1 silenced cells displayed a greater than 2-fold reduction in their invasion capacity in hypoxic culture conditions (**Fig. 6A**). The difference in the invasion capacity between control and knockdown cells was far more dramatic in the hypoxic condition than in the normoxia culture (**Fig. 5G** and **Supplementary fig. S6A**). These observations indicated that LOXHD1 may play a role in EwS cell response to hypoxia. Therefore to better understand this, using RNA-seq, we studied hypoxia-induced transcriptome changes in parental and LOXHD1 silenced SK-N-MC cells (**Fig. 6B**). Using differential expression analysis (36) for the hypoxic samples, we first identified 204 genes with > 4-fold upregulation and 77 genes with > 4-fold downregulation (p-value <0.001), suggesting a robust transcriptional response to hypoxia in the parental SK-N-MC cells (**Fig. 6C**). As expected, GSEA for the altered transcriptome showed strong positive enrichment (NES=2.97) for the hallmark Hypoxia signature (**Supplementary fig. S6B**). The Gene Ontology (GO) for the 204 upregulated genes comprised processes and pathways associated with hypoxic response and HIF-1 signaling (**Supplementary fig. S6C**), confirming that the SK-N-MC cells are sensitive to hypoxia. We next evaluated the role of LOXHD1 in EwS hypoxic response in the eKD1 and eKD2 cells. Unlike the control cells, these two independent LOXHD1 silenced clones showed weaker hypoxic stress response. GSEA analysis showed a significant reversal of both the upregulated and downregulated gene signature (**Fig. 6D**), suggesting a weakened hypoxia response in LOXHD1 silenced EwS cells. HIF1α is the main transcription factor involved in the transcriptional response to hypoxia (37). Under hypoxic conditions, HIF1α is primarily stabilized by functional inactivation of prolyl hydroxylases (PHDs) and VHL E3 ligase complex, which label HIF1α for proteasomal degradation (38). Remarkably, hypoxia-treated eKD1 and eKD2 SK-N-MC cells showed less HIF1α protein levels than parental control cells (**Fig. 6E**) despite a lack of downregulation of its mRNA expression (**Supplementary fig. S6D**). There was a slight increase in *HIF1A* mRNA expression in the hypoxia treated eKD cells compared to that of the control cells, which could potentially be a result of compensation to restore HIF1α protein in these cells. This data suggests that LOXHD1 silencing most likely destabilize HIF1α protein rather than reduce the transcription of *HIF1A*. The coordinated activity of iron-dependent PHDs maintains the appropriate balance of HIF1α protein, and iron chelators such as desferrioxamine (DFO) result in HIF1α stabilization (39,40). However, treatment with DFO did not result in differential stability of HIF1α in eKD cells compared to controls, ruling out a potential deficiency of canonical HIF1α regulatory signaling in the LOXHD1 silenced cells (**Supplementary fig. S6E**). To determine if LOXHD1 is directly involved in HIF1α regulation, we carried out co-immunoprecipitation experiments following ectopic expression of HIF1A and LOXHD1. Co-transfection of HA-tagged HIF1α and Myc-tagged LOXHD1 in 293T fibroblasts grown under hypoxic condition, followed by immunoprecipitation with MYC-tag LOXHD1 was able to pulldown HA-tagged HIF1α (**Fig. 6F**), suggesting a direct physical interaction between these two proteins. Altogether, these results demonstrate that LOXHD1 functions as a regulator of HIF1α stability and its transcriptional activity in EwS cells.

**Figure 6:**
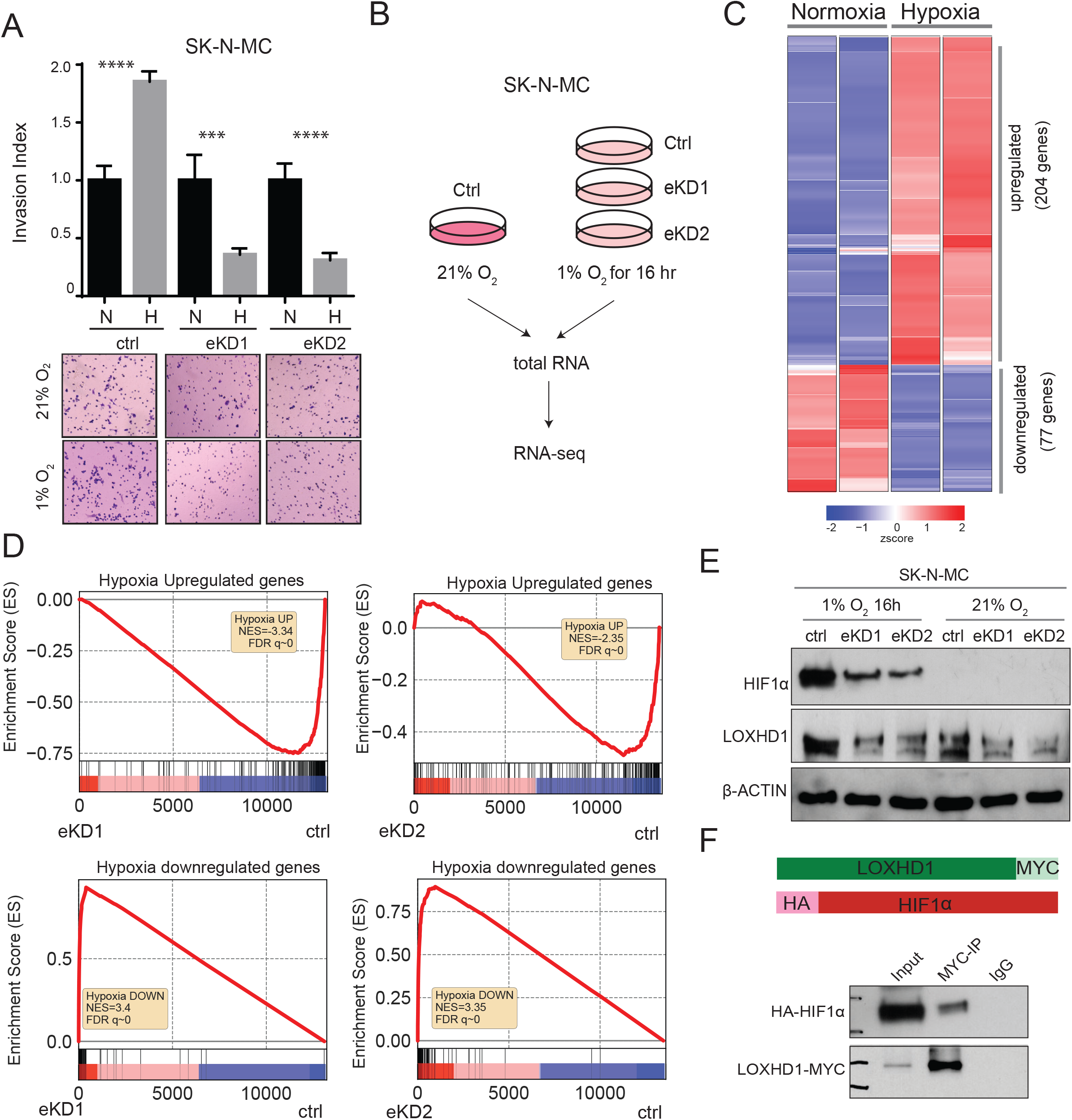
LOXHD1 silencing impairs EwS response to hypoxia. **(A)** Hypoxia induces impaired invasion response in LOXHD1 knockdown cells. *Top* Quantification of a Boyden chamber Matrigel invasion assay performed with control, eKD1 and eKD2 cells at 21% O_2_ and 1% O_2_ for 24h. *bottom*, representative images of each indicated group. (Quantification of the hypoxia only condition is in Supplementary Fig 6a.). **(B)** Schematic showing the hypoxia RNA-seq experimental design. **(C)** Hypoxia induces major transcriptional changes in LOXHD1 intact EwS cells. Heatmap shows differential expressed genes under hypoxia (1% O_2_ for 16h) compared to normoxia in SK-N-MC cells. **(D)** RNA-seq followed by GSEA showing negative enrichment of the hypoxia upregulated signature and positive enrichments of the hypoxia downregulated signature in *LOXHD1* eKD1 and eKD2 SK-N-MC cells. **(E)** LOXHD1 silencing reduces HIF1α stabilization under hypoxia. Immunoblots of HIF1α and LOXHD1 in control and eKD cells under hypoxia and normoxia culture. β-actin used as loading control. **(F)** LOXHD1 interacts with HIF1α. HEK293T cells were co-transfected with MYC-tagged LOXHD1 and HA-tagged HIF1α, and cultured under hypoxia for 24h. Total protein lysates used for immunoprecipitation with MYC-tag antibody. *top*, Schematic showing the structure of the two constructs. *bottom*, Immunoblots of anti-HA and anti-Myc antibodies.

### LOXHD1 silencing affects EwS metastasis and tumor growth *in vivo*

Finally, to examine the role of LOXHD1 in the EwS tumor growth and metastasis *in vivo*, we employed three different metazoan models such as the chicken chorioallantoic membrane (CAM) model, zebrafish model, and mouse xenograft model. In the CAM assay, cancer cells introduced on the upper CAM-proliferate, invade the basement membrane, intravasate the nearby vasculature, and circulate in the blood vessels that can be captured at the lower CAM, thereby providing an estimate of their invasion and intravasation potential (**Fig. 7A**) (41–43). We observed a significantly impaired invasion and intravasation to the lower CAM by the enhancer silenced LOXHD1 depleted SK-N-MC and RD-ES cells than their respective parental controls (**Fig. 7B**). We next studied EwS metastasis in a zebrafish model, which has been widely used to test the metastatic potential of various human cancer cell lines, including EwS (35,44). Strikingly, the zebrafish embryos displayed a significantly impaired metastatic dissemination of RFP labeled LOXHD1 depleted RD-ES cells from the yolk sac to the tails and head of the embryos, than the parental controls, providing strong evidence for LOXHD1 as a mediator of EwS metastasis *in vivo* (**Fig. 7C** and **7D**). We next tested the effect of LOXHD1 silencing by CRISPRi in the murine SK-N-MC xenograft model. Compared to parental control, the LOXHD1 silenced SK-N-MC xenograft demonstrated a significantly reduced tumor growth (**Fig. 7E** and **7F**), which was accompanied by increased necrotic margins and lower mitotic foci (**Supplementary fig. S7A-B**). This result is in agreement with the colony and sphere formation assay performed *in vitro*, and together provides concrete evidence supporting that LOXHD1 promotes EwS tumorigenicity. This short-term subcutaneous xenograft assay may not be an ideal model to study EwS spontaneous metastasis, and tail vein injection experiment can only test the colonization ability of the tumor cells. Together, these *in vivo* data clearly establish the role of LOXHD1 in regulating the EwS tumor formation and metastasis (**Fig. 7G**).

**Fig. 7:**
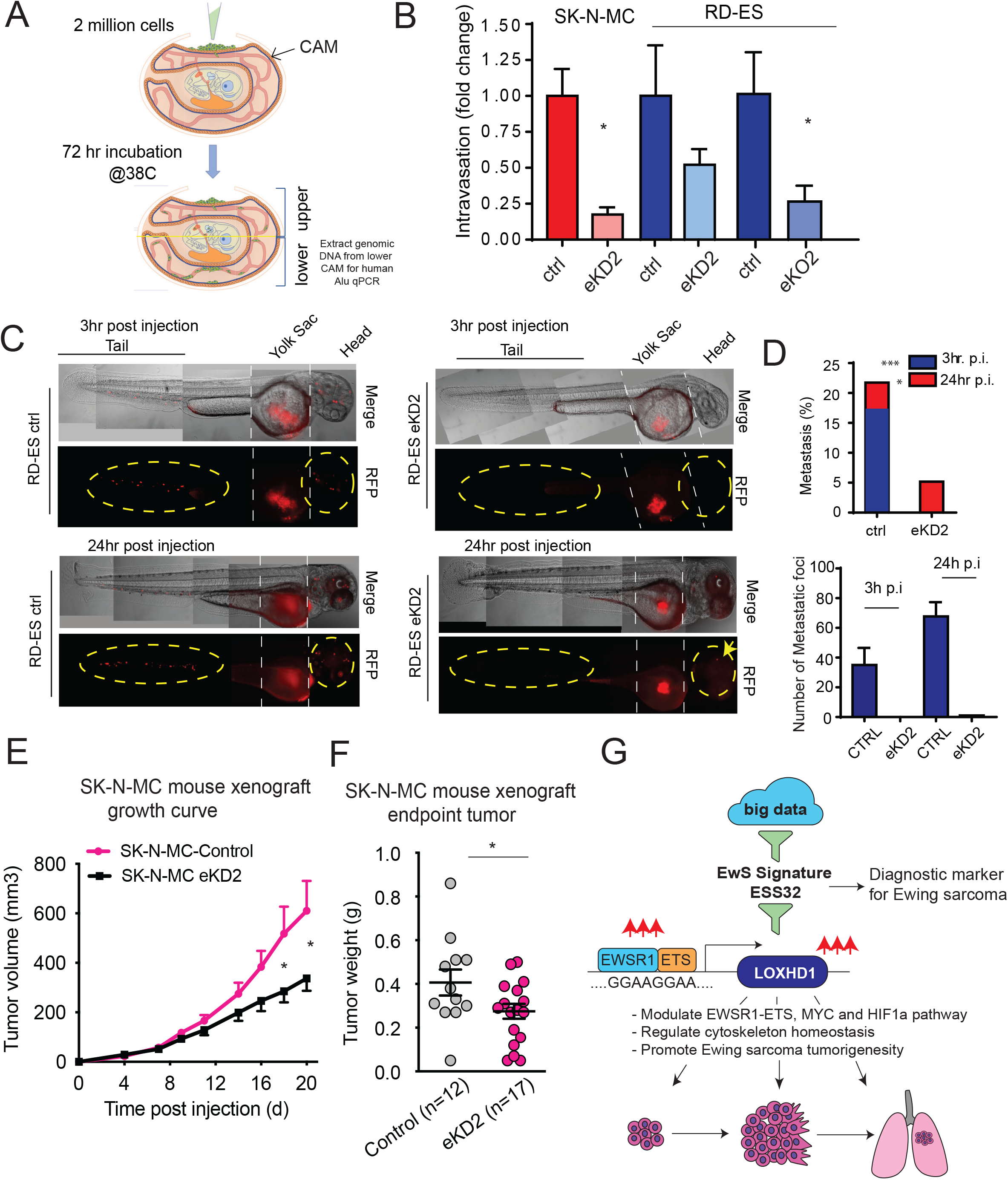
LOXHD1 silencing attenuates the oncogenic and metastatic phenotype of EwS cells *in vivo*. LOXHD1 knockdown reduce cell intravasation in a chicken CAM model. **(A)** Schematic showing CAM intravasation assay. Two million cells are cultured atop the embryonic chick upper CAM for 3 days followed by genomic DNA isolation from the lower CAM, which is used to measure the intravasated human cells by qPCR using human-specific Alu primers. **(B)** Bar graph showing normalized fold difference in the intravasated cells for the indicated group. *p<0.01, by students t-test. **(C)** LOXHD1 knockdown impairs EwS metastasis in a zebrafish model. Representative images of zebrafish in the control and LOXHD1 knockdown group showing metastasis in yellow circles and arrow at 3h and 24h post-injection. **(D)** *Top* Bar graph of percentages of zebrafish harboring metastasis at 3h and 24h time point. *Bottom* Quantification of total number of metastatic foci. ***p<0.001, *p<0.01 by chi-square test. **(E)** LOXHD1 silencing attenuates tumor formation in mice. Growth curve of the xenograft experiment using SK-N-MC control and isogenic eKD2 cells. **(F)** Bar graph of the endpoint tumor weights. **(G)** Schematic illustrating the discovery and the role of LOXHD1 in influencing multiple oncogenic pathways in Ewing sarcoma genesis and progression. **p<0.01, by Students t-test.

## Discussion

In this study, we have identified the stereociliary protein LOXHD1 as a highly specific EwS gene product with oncogenic and metastasis promoting properties. Our results demonstrate the EWSR1-FLI1 mediated *de novo* enhancer activates the expression of this developmentally silenced gene. While previous work has established LOXHD1 mutation in DFNB77, a progressive form of autosomal-recessive nonsyndromic hearing loss (ARNSHL) (23), we provide the first evidence of its role in cellular physiology and, in particular, EwS tumorigenicity. Additionally, through integrative transcriptomic analysis, the identification of ESS32 gene signature comprising known and new EWSR1-FLI1 targets which accurately stratify metastatic EwS from non-EwS samples is a critical discovery with translational potential as a EwS diagnostic and prognostic biomarker.

Besides LOXHD1, there are close to 20 genes in the human genome that code for proteins containing the PLAT domain, including ALOX12, LPL, PKD1, and RAB6IP1. Except for LOXHD1 with multiple PLAT domains, other proteins possess a single PLAT domain. The highly conserved PLAT domain is involved in protein-protein and protein-lipid interactions *(45)*. Usually, they tend to associate peripherally with the cytosolic side of the plasma membrane and mediate interactions with other transmembrane signaling proteins (24). Interestingly, LOXHD1 is required for hearing in human and mice, and localizes between the membrane and the actin-cytoskeleton of stereocilia, potentially to connect them (23). Further work will be needed to dissect the exact contribution of the multiple PLAT domains of LOXHD1 in protein-protein and lipid-protein signaling, but our work highlights its potential role in cytoskeleton organization. In addition to the 13 PLAT domains, we found a coiled-coil domain in LOXHD1 that has not been characterized previously (23). Coiled-coil domain containing proteins are associated with critical biological functions such as transcription and cell movement. Notable examples are the transcription factor c-Fos and c-Jun, as well as the muscle protein tropomyosin (46). Further, the identification of nuclear localization signal (NLS) near the coiled-coil domain in LOXHD1, its localization to the nucleus, its effect on EWSR1-FLI1, MYC transcriptional program, and its potential role in HIF1a stability under hypoxia suggests a direct role of this enigmatic protein as a mediator of oncogenic functions in EwS cells. Further research is ongoing to delineate the molecular mechanism of LOXHD1 mediated cytoskeleton organization, transcriptional regulation and hypoxic stress response in EwS cells.

Oncofusions as driver oncogenes are particularly common in pediatric cancer. Some fusions provide new therapeutic targets. For instance, both BRAF inhibitors and MEK inhibitors have been tested with limited success in pilocytic astrocytoma bearing BRAF fusions (47–49). Alveolar soft part sarcoma and a subset of renal cell carcinoma patients with TFE3 fusions are more likely to respond to MET inhibitors (50). However, small molecule-based targeted therapies toward EWSR1-FLI1 and associated proteins have not been successful in EwS. Immunotherapy has emerged as the next frontier in cancer treatment (51). Tumors with high mutation load often generate T cells against neoantigens derived from mutated proteins. In these settings, immune checkpoint blockade can lead to therapeutic responses (52). Sarcomas are extremely diverse with >50 diagnostic subtypes, most of which (like EwS) have low mutation load (53,54). Response rates to anti-CTLA-4 or anti-PD-1 single-agent treatment are limited and appear to be restricted to specific histologic subtypes such as undifferentiated pleomorphic sarcoma, liposarcoma, leiomyosarcoma, and synovial sarcoma (55). Adoptive cell therapy (ACT) with T cells engineered to recognize non-mutated tumor-associated antigens offers an attractive alternative. This is supported by encouraging clinical trial results with TCR gene therapy directed against the NY-ESO-1 tumor antigen in patients with synovial sarcoma and metastatic melanoma demonstrating a durable complete cancer regression (56,57). These results have stimulated efforts to genetically modify lymphocytes to improve their specific antitumor efficacy and to extend the range of tumors that can be targeted. However, a significant impediment to the development of effective immune-based therapies for EwS is in identifying tumor-specific molecules with a limited expression in healthy tissues. Ideally, the target antigen has to be derived from a protein that is 1-highly expressed in tumor cell (to ensure on-target activity), 2-minimally expressed in normal tissue (to reduce off-target activity/toxicity), and 3-required for tumor cell survival/sustenance (to prevent therapy resistance) (58–60). Our observations demonstrating the highly exclusive expression pattern of LOXHD1 and functional validation of its oncogenic potential fulfill these criteria for a potential ACT-based immunotherapy against LOXHD1 in EwS.

In summary, our findings identify LOXHD1, which is transcriptionally silent in the vast majority of normal and cancer cells, as a direct EWSR1-FLI1 target gene that plays an important role in cytoskeletal homeostasis and oncogenic transcription in EwS. We show that loss of LOXHD1 expression through deletion or epigenetic silencing using dCas9-KRAB of its upstream EWSR1-FLI1 bound GGAA microsatellite *de novo* enhancer strongly inhibits the tumorigenic potential of EwS cells *in vitro* and *in vivo*. While there is undoubtedly more functional characterization of LOXHD1 needs to be made, we believe this study provides a strong basis for identifying LOXHD1-derived endogenous peptide epitopes in EwS cells for ACT-based immunotherapy for this deadly disease.

## Acknowledgments

We thank Dr. Frank S. Lee for HIF1α plasmid and Dr. Terrance Gades for providing technical assistance with the hypoxia chamber. We thank Dr. Arul Chinnaiyan and Dr. Gerald Linette for insightful discussion. This work was partly funded by Hayman Family foundation; and Sarcoma Foundation of America Research Award to I.A.A. Research in I.A.A laboratory is also supported by NIH (1-R01 CA249210-0), Department of Defense Idea Development Award (W81XWH-17-0404) to I.A.A. The laboratory of T.G.P.G. was supported by the Dr. Leopold and Carmen Ellinger Foundation, the Dr. Rolf M. Schwiete Foundation, the German Cancer Aid (DKH-70112257 and DKH-70114111), the Gert und Susanna Mayer Foundation, the SMARCB1 association, and the Barbara und Wilfried Mohr Foundation.

## Contributions

I.A.A. conceptualized the study. Q.D. conducted the experiments with help from S.A., R.R., P.W., Z.C. R.N. performed all the bioinformatics analysis. F.C-A. and T.G.P.G. designed and performed mouse xenograft experiment. M.C., C.K-S., K.W., T.S.K.E-M., N.G., and T.G., provided reagents, expertise and feedback. Q.D., R.N., and I.A.A. wrote the manuscript with input from all the authors.

## Data Deposits

The RNA-seq and ChIP-seq data have been deposited in the NCBI GEO database with the accession number GSE163335 (Token: oxiziyamlhsjlqj)

