## supplementary figures and methods for "Oncofusion-driven *de novo* enhancer assembly promotes malignancy in Ewing sarcoma *via* aberrant expression of the stereociliary protein LOXHD1"

### Supplementary Figure 1

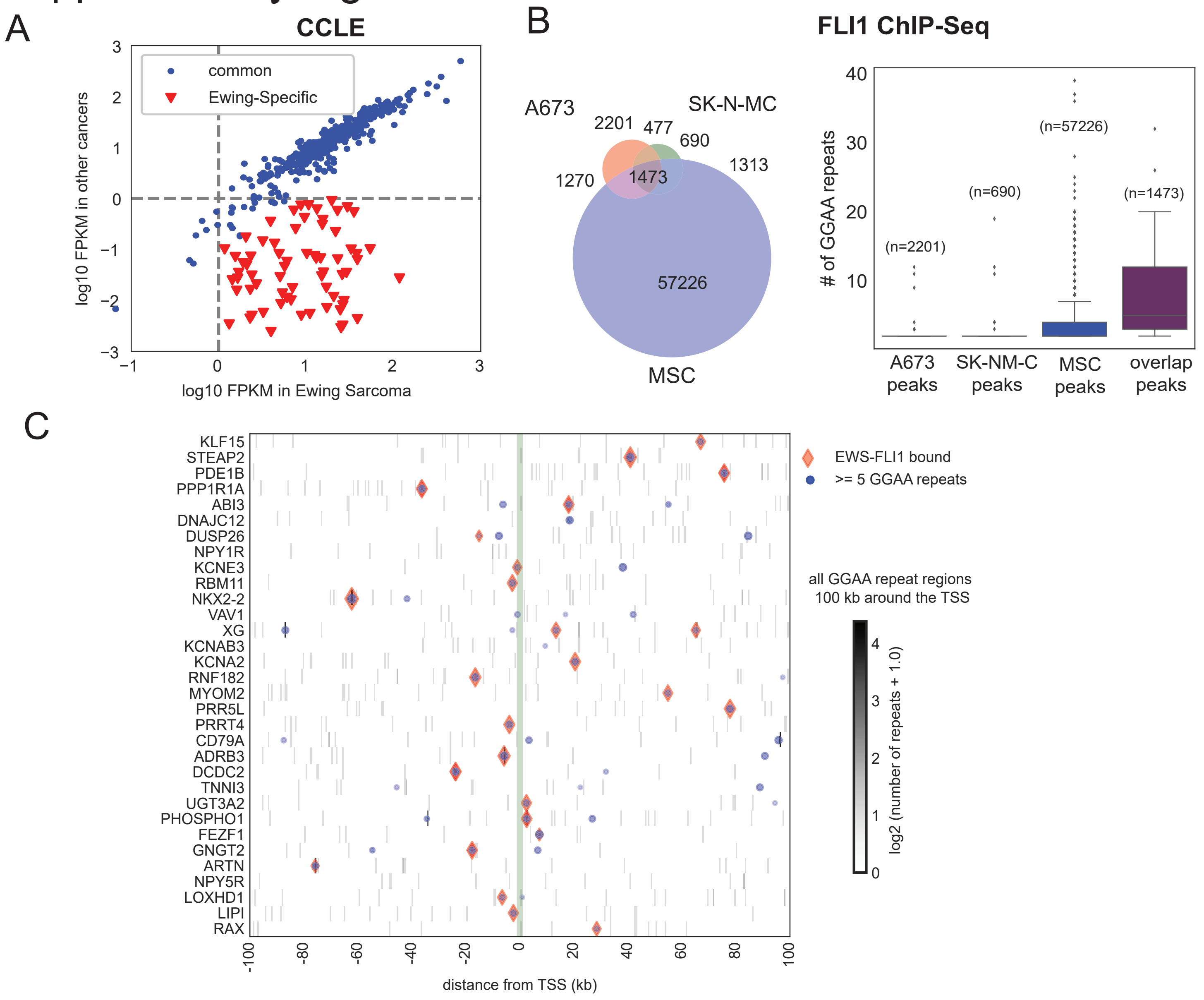


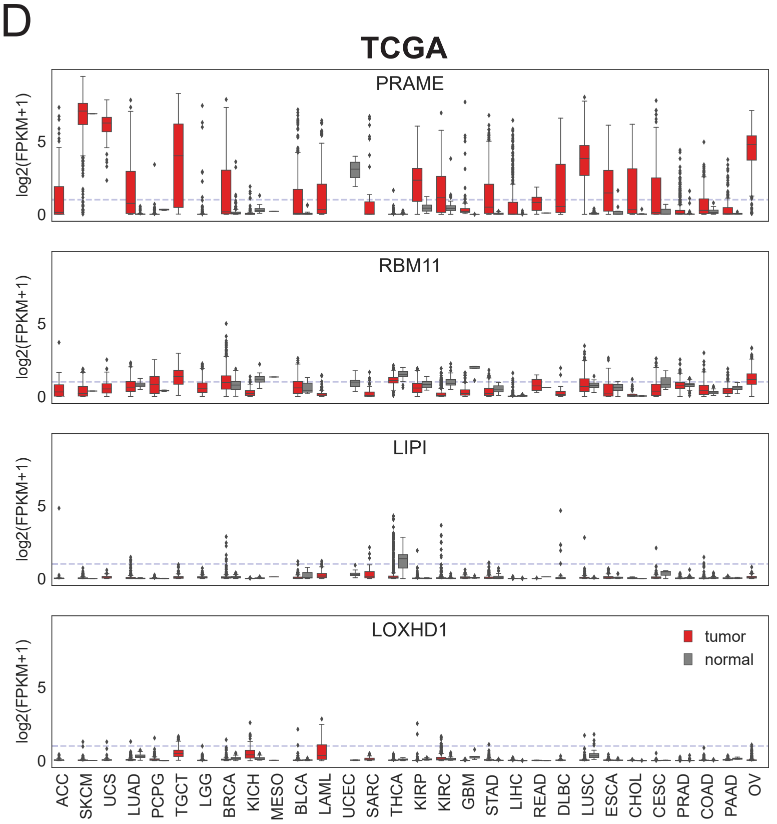


| code | cancer | normal | tumor |
|---|---|---|---|
| LAML | Acute Myeloid Leukemia | 0 | 151 |
| ACC | Adrenocortical carcinoma | 0 | 79 |
| BLCA | Bladder Urothelial Carcinoma | 19 | 411 |
| LGG | Brain Lower Grade Glioma | 0 | 529 |
| COAD | Colon adenocarcinoma | 41 | 464 |
| ESCA | Esophageal carcinoma | 11 | 162 |
| GBM | Glioblastoma multiforme | 5 | 168 |
| KICH | Kidney Chromophobe | 24 | 65 |
| KIRP | Kidney renal papillary cell carcinoma | 32 | 289 |
| LIHC | Liver hepatocellular carcinoma | 50 | 374 |
| LUAD | Lung adenocarcinoma | 59 | 526 |
| LUSC | Lung squamous cell carcinoma | 49 | 501 |
| DLBC | Lymphoid Neoplasm Diffuse Large B-cell Lyr | 0 | 42 |
| MESO | Mesothelioma | 0 | 1 |
| OV | Ovarian serous cystadenocarcinoma | 0 | 379 |
| PAAD | Pancreatic adenocarcinoma | 4 | 178 |
| PCPG | Pheochromocytoma and Paraganglioma | 3 | 150 |
| PRAD | Prostate adenocarcinoma | 52 | 499 |
| READ | Rectum adenocarcinoma | 1 | 6 |
| SARC | Sarcoma | 0 | 35 |
| SKCM | Skin Cutaneous Melanoma | 1 | 471 |
| STAD | Stomach adenocarcinoma | 32 | 375 |
| TGCT | Testicular Germ Cell Tumors | 0 | 156 |
| THCA | Thyroid carcinoma | 58 | 510 |
| UCS | Uterine Carcinosarcoma | 0 | 56 |

### Supplementary Figure 1 (contd)

E

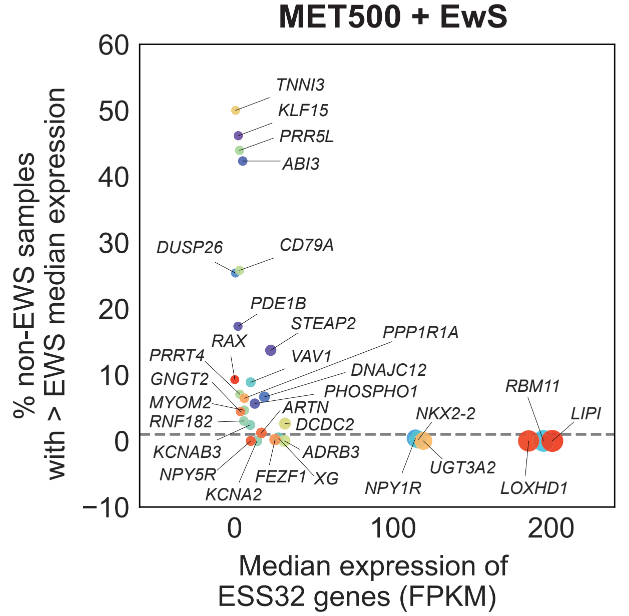


F

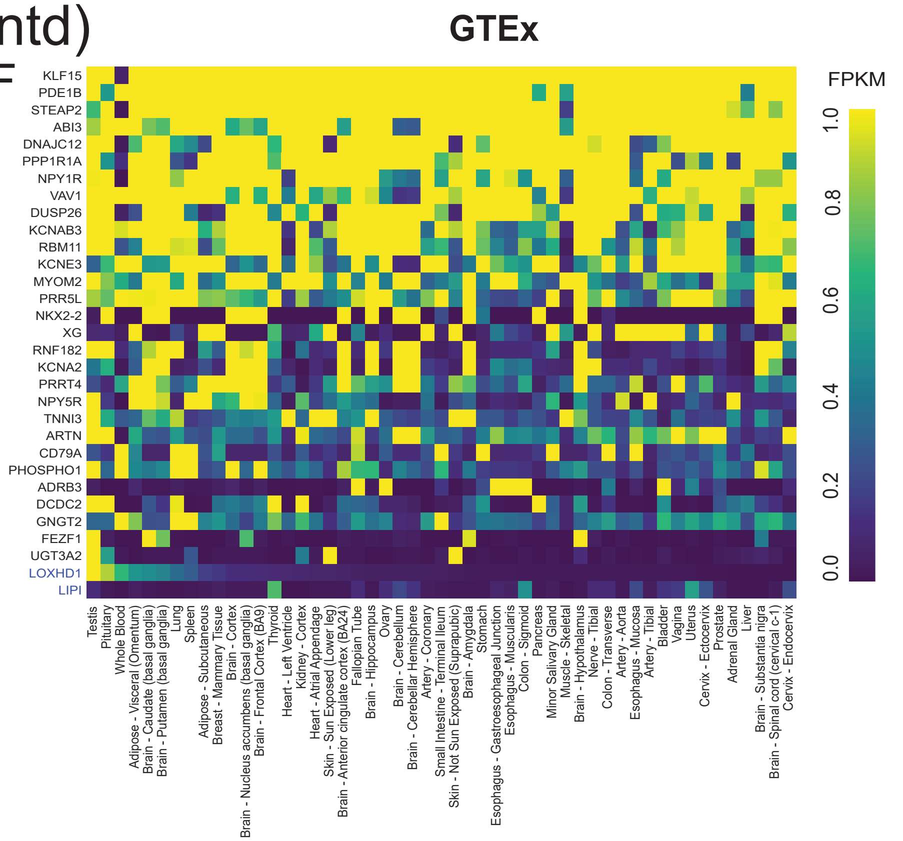


G

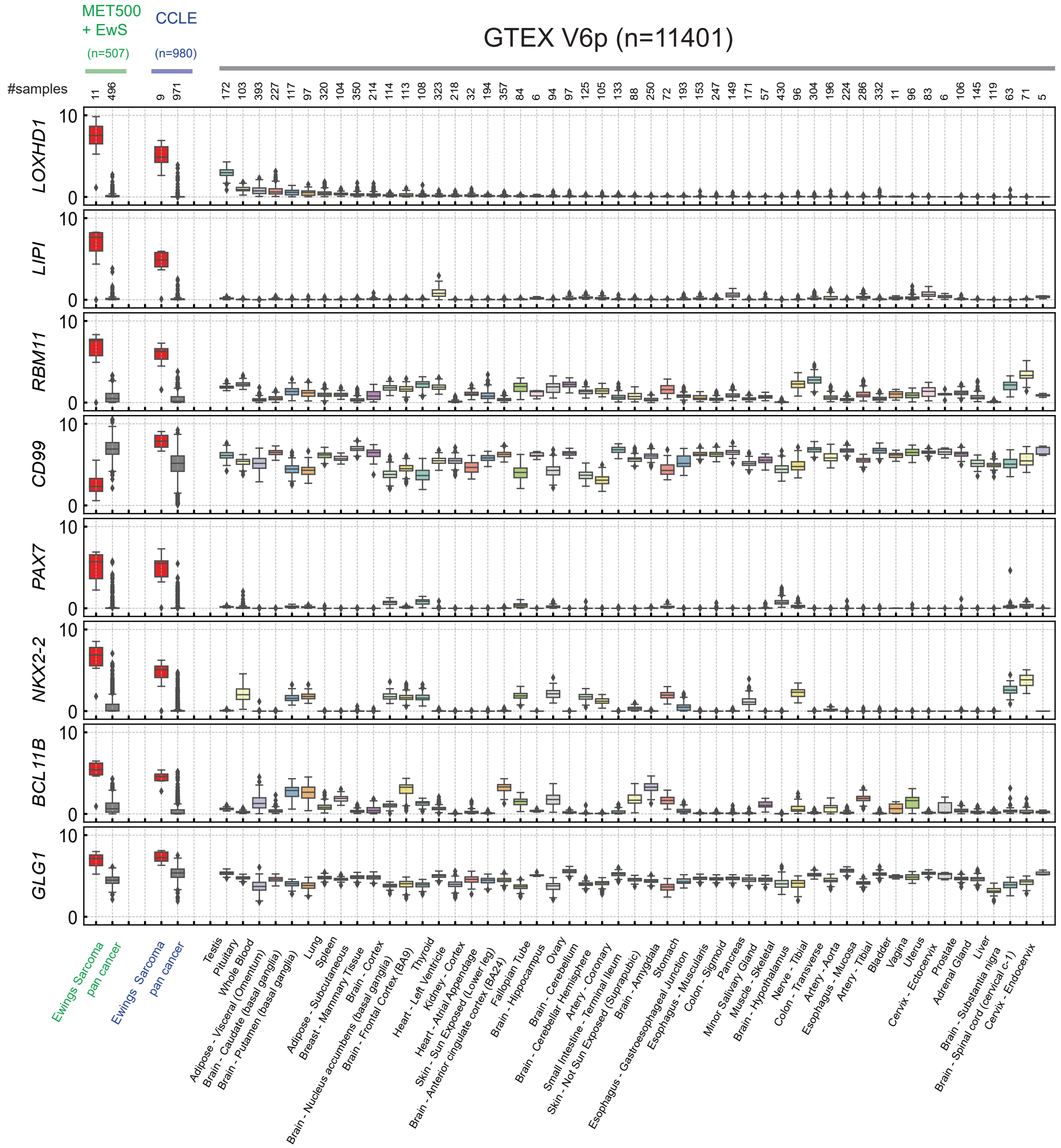


H

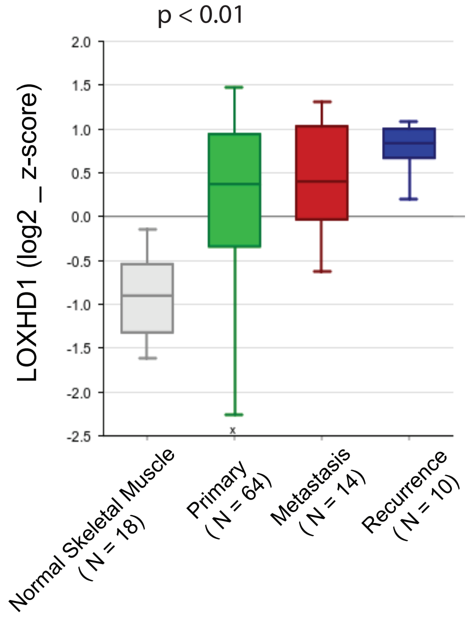


I

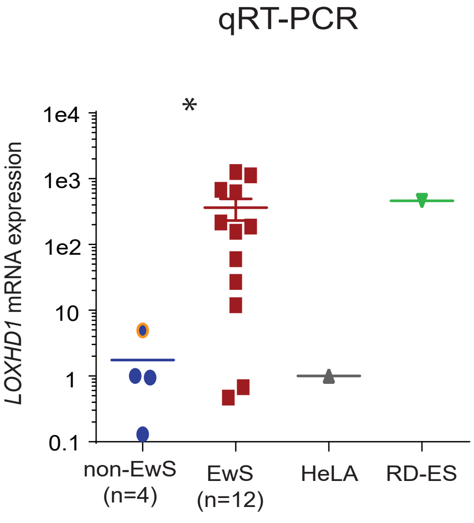

**Supplementary Figure 1: Integrative analysis of ChIP-seq and CCLE, MET500, TCGA and GTEx transcriptomic datasets.** **(A)** *Analysis of CCLE dataset;* Expression of the 516 genes (step 1 of Fig. 1a) in EwS vs other cell lines. The 89 EwS specific genes, seen in the lower-right quadrant and encircled, are expressed only in EwS cell lines and show <1 FPKM expression in all others. **(B)** *Analysis of FLI1-ChIP-seq data; (left)* Overlap analysis between FLI1 enriched ChIP-Seq peaks in A673 (GSM1517562), SK-NM-C (GSM1517537), and EWSR1-FLI1 overexpressed MSCs (GSM2472088, GSM2472102, GSM2472108) yields 1473 conserved FLI1-bound regions, *(right)* distribution of the number of GGAA microsatellite repeats contained within A673-specific, SK-N-MC-specific, MSC-specific and the 1473 overlapping regions shows pronounced enrichment of the GGAA microsatellites in the overlapping regions. **(C)** Colormap shows the locations of GGAA microsatellite repeat regions $\pm$100 kb around the TSS of the ESS32 genes. Symbol overlays represent regions with >= 5 GGAA repeats (blue circles) and contain a FLI1 ChIP-seq enrichment peak (red diamonds). **(D)** *Analysis of TCGA dataset;* Boxplot showing nearly zero expression of *LOXHD1* and *LIPI* in all TCGA cancer-subtypes in tumor and normal samples. The expression of *RBM11* located in *LIPI* locus is shown. . Expression of PRAME is shown as an example of gene that is expressed across all cancer-subtypes. Abbreviation of codes for cancer-subtypes and the corresponding numbers of normal and tumor samples are shown alongside. **(E)** *Analysis of MET500 RNA-seq dataset;* Plot showing the median expression of the ESS32 genes (step 3 in Fig. 1a) in EwS against the percentage of non-Ewing samples that show expression higher than the median value in EwS. The dotted line marks the cutoff of 1%. **(F)** *Analysis of GTEx datasets;* Heatmap showing the expression of the ESS32 genes showing null expression of *LOXHD1*, *LIPI* in all tissues except testis. **(G)** Box plots comparing the expressions of indicated targets across MET500 + EwS (n=507), CCLE (n= 980), and GTEX (n=11401) samples. (**H**) Box plots showing LOXHD1 expression in the indicated samples. Publicly available Affymetrix dataset GSE34620 comprising 117 EwS samples was analyzed and Log2 z-score is plotted. $p < 0.01$ One-way Anova between the groups. **(I)** qRT-PCR analysis of *LOXHD1* mRNA expression in an independent cohort of EwS tumor tissues, non-EwS tissues (testis is indicated with orange border), HeLa and RD-ES cells were used as negative and positive control, respectively. * $p < 0.001$, by two-tailed Student's t test.

### Supplementary Figure 2

A

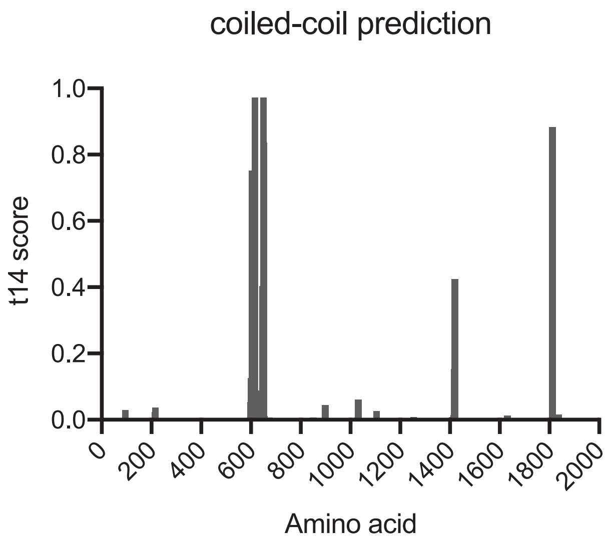


B

| Predicted monopartite NLS | | |
|---|---|---|
| **Pos.** | **Sequence** | **Score** |
| 616 | LQRKKKKRKGSDE | 15 |
| 617 | QRKKKKRKGS | 6 |
| 618 | RKKKKRKGSD | 7 |
| 618 | RKKKKRKGSDE | 6 |
| 1741 | IPLKRKRKYFKVF | 6 |
| 1741 | IPLKRKRKY | 7 |
| 1742 | PLKRKRKYFKV | 8.5 |
| 1743 | LKRKRKYFKVF | 8.5 |

C

LOXHD1- NLS coiled coil domain

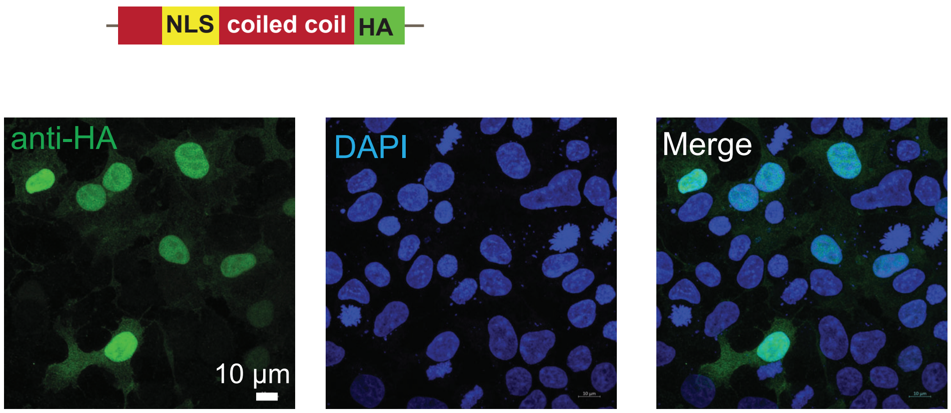

**Supplementary Figure 2: Domain prediction of LOXHD1 protein, and the effect of LOXHD1 overexpression in Hela cells. (A)** Coiled-coil structure prediction by COILS (https://embnet.vital-it.ch/software/COILS_form.html), y-axis shows the probability of a 14 amino acid (aa) coiled-coil structure and the x-axis shows the amino acids position of LOXHD1 protein. The predicted coil-coil structure is seen as the peak between aa 653 to aa 677. **(B)** Nuclear localization signals (NLS) predicted by cNLS Mapper (http://nls-mapper.iab.keio.ac.jp/cgi-bin/NLS_Mapper_form.cgi), NLS with a score larger than 8 is exclusively localized to the nucleus. **(C)** Functional validation of NLS. Immunofluorescence staining with HA antibody showing nuclear signal in HEK293T cells transfected with plasmid containing HA-tagged NLS-coiled-coil domain of LOXHD1.

Supplementary Figure 3

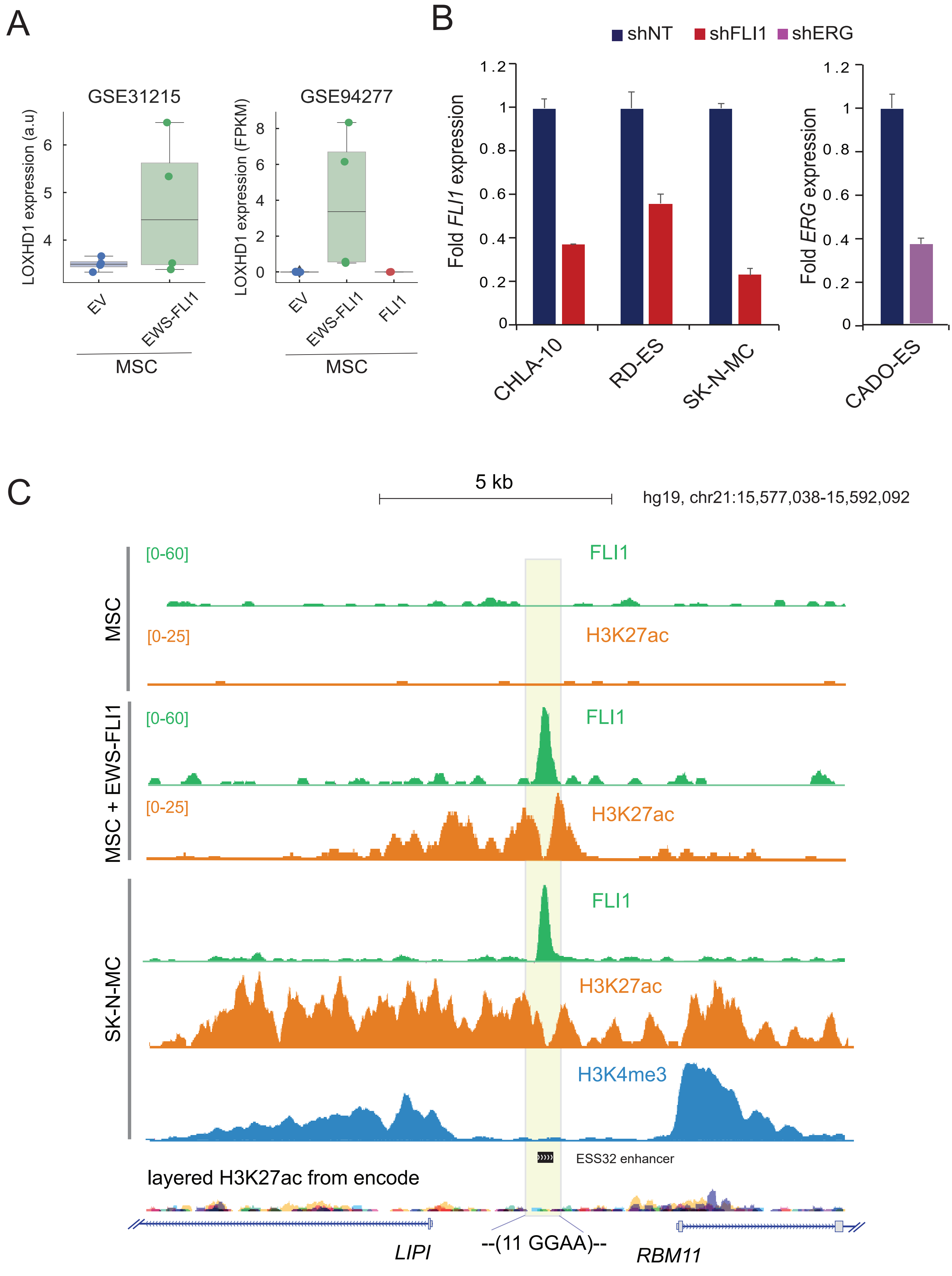

**Supplementary Figure 3: *LOXHD1* expression in EWSR1-FLI1 overexpressing MSCs and FLI knockdown EwS cell lines. (A)** *(left)* Comparison of *LOXHD1* expression profiled using Affymetrix Human Genome U133 Plus 2.0 Array (GSE31215) in MSCs expressing EWSR1-FLI1 (n=4), *(right)* Comparison of *LOXHD1* FPKM expression in MSCs expressing EWSR1-FLI1 (n=4) and wild type FLI1 (n=4), profiled using RNA-seq (GSE94277). **(B)** Bar graph showing qRT-PCR results of FLI1 (*left*) and ERG (*right*) expression with shRNA knockdown of EWSR1-FLI1 and EWSR1-ERG respectively in the indicated EwS cells. **(C)** Genome browser view showing the de novo enhancer assembly in the vicinity of *LIPI* and *RBM11* locus in EWSR1-FLI1 overexpressing MSCs and SK-N-MC cells.

### Supplementary Figure4

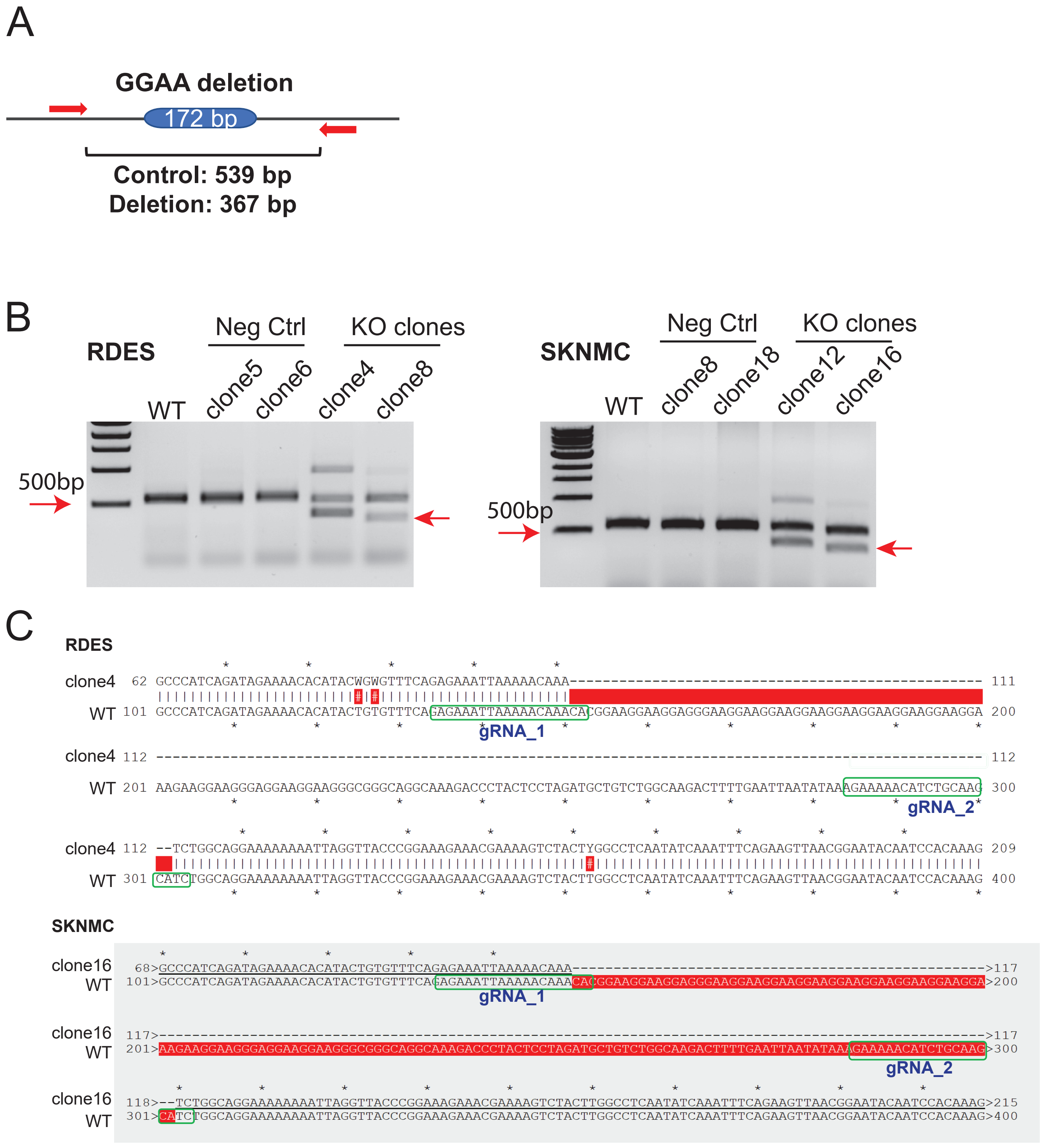

**Supplementary Figure 4: Verifications of the LOXHD1 enhancer knockout single clones. (A)** PCR and sequencing strategy for GGAA microsatellite deletion. Primers designed outside of the two gRNAs flanking the GGAA microsatellite; wild-type allele predicted to have 539bp amplicon while the deleted allele is predicted to have 367bp amplicon. **(B)** DNA gel image of PCR products with the genomic DNA of empty cas9 control cells and the enhancer targeting sgRNA CRISPR-cas9 virus infected cells. SK-N-MC and RD-ES cells were transduced with empty gRNA cas9 virus or cas9 with two gRNAs flanking the GGAA microsatellite, three days post viral transduction cells were subjected for puromycin selection for three days and harvested for genomic DNA for PCR. Arrow indicates the allele with 172bp deletion. **(C)** Sanger sequencing result of the enhancer knockout RD-ES and SK-N-MC single cell derived clones. Red region shows the deletion between two guide RNAs.

### Supplemetnary Figure 5

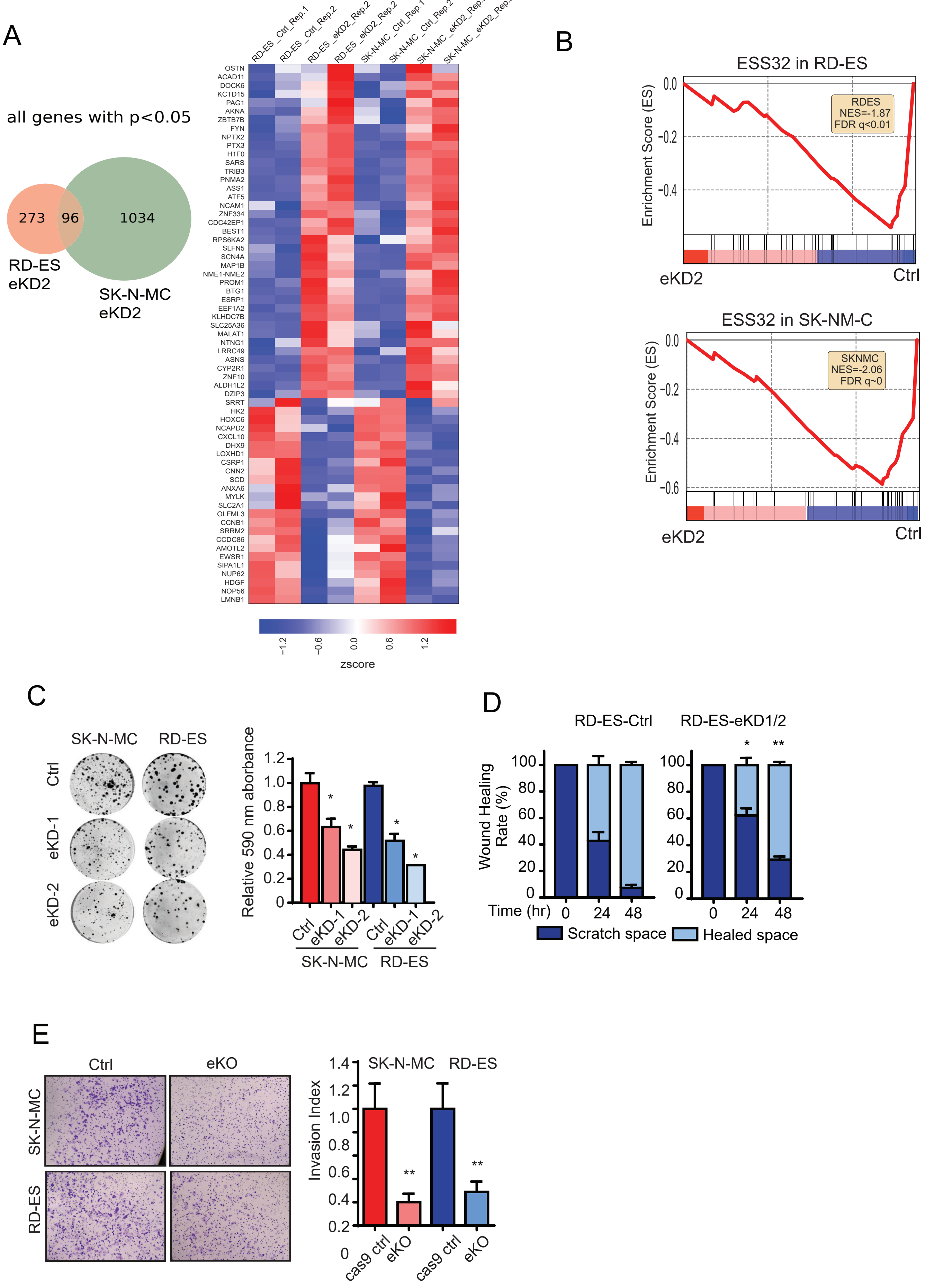

**Supplementary Figure 5: LOXHD1 loss impairs EwS signature and tumor phenotype in vitro. (A)** Venn diagram and heatmap showing differentially expressed genes in LOXHD1 enhancer knockdown (eKD2) cells. **(B)** RNA-seq followed by GSEA showing negative enrichment of the ESS32 signature in *LOXHD1* eKD2 RD-ES and SK-N-MC cells. LOXHD1 knockdown impairs colony formation ability in EwS cells. **(C)** *left* Representative images of a colony formation assay performed by seeding 5000 control and eKD cells in each well of 6-well plates, each group is done with triplicates. *right*, bar graph showing quantifications of the crystal violet staining of the colonies. LOXHD1 knockdown slows down cell migration in EwS cells. **(D)** Bar graph of a wound healing assay presented by quantifications of the scratch widths in RD-ES control and eKD cells. LOXHD1 knockdown reduces cell invasion in EwS cells. **(E)** *left*, representative images of a Boyden chamber invasion assay with SK-N-MC and RD-ES control and eKO single cell clones. *right*, bar graph showing quantifications of the invaded cells.

### Supplementary Figure 6

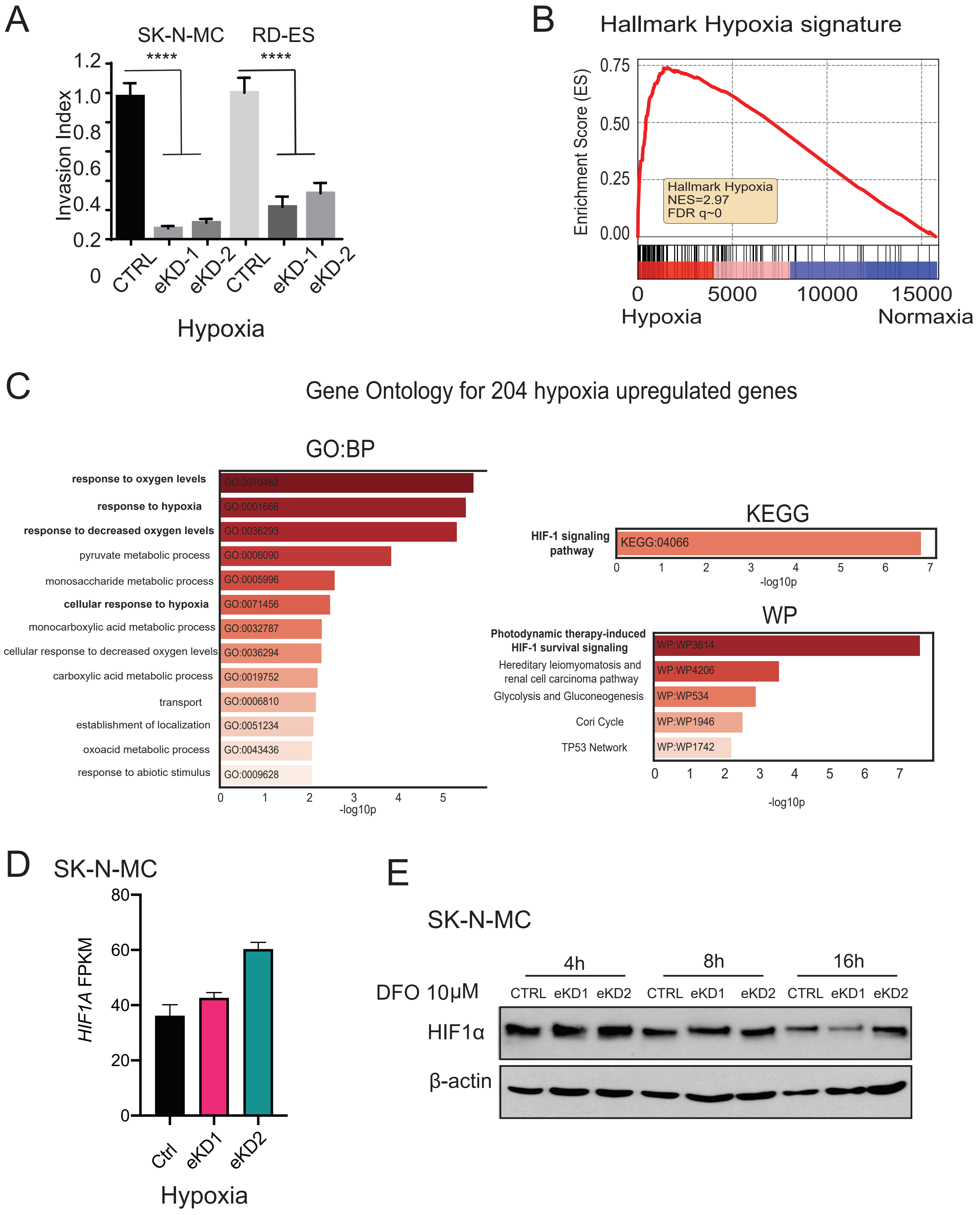

**Supplementary Figure 6: LOXHD1 knockdown diminishes hypoxic responses in EwS cells.** Hypoxia enhances the effects of LOXHD1 knockdown on EwS cell invasion. **(A)** Bar graph showing quantifications of invasion index of the hypoxia samples in the experiment of Fig. 6a. LOXHD1 proficient SK-N-MC cells present strong hypoxia response. (**B)** RNA-seq followed by GSEA showing a positive enrichment of Hallmark Hypoxia signature in the hypoxic treated control SK-N-MC cells compared to the normoxia sample. **(C)** Gene Ontology for the 204 genes induces by hypoxia showing hypoxic responses and HIF-1signaling among the top of the lists. LOXHD1 knockdown in EwS cells does not affect *HIF1A* transcription. **(D)** Bar graph of the FPKM of *HIF1A* in control and two eKD SK-N-MC cells under hypoxia. **(E)** Immunoblot of HIF1$\alpha$ with DFO treatment at 10 $\mu$M for 4, 8 and 12h in control and two eKD SK-N-MC cells. $\beta$ actin was used as loading control.

### Supplementary Figure 7

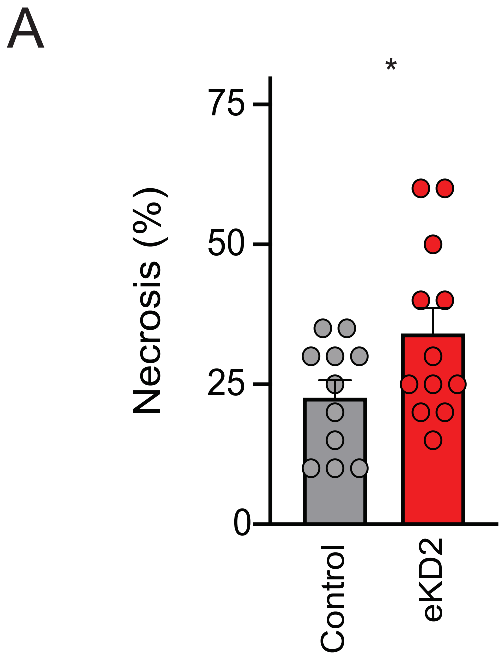


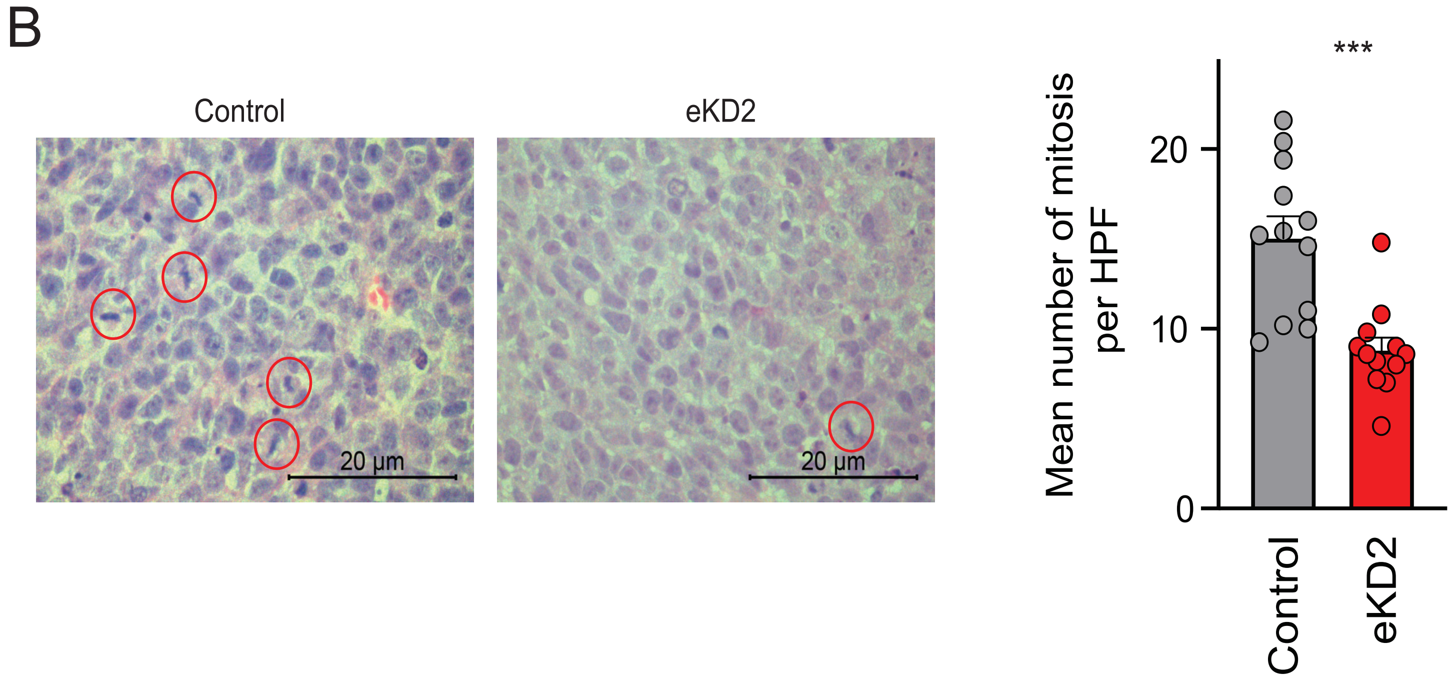

**Supplementary Figure 7: LOXHD1 knockdown affects EwS growth in vivo. (A)** Bar graph showing the percentage of necrotic regions in the control and LOXHD1 enhancer knockdown SK-N-MC xenograft. **(B)** Representative images showing mitotic nuclei in the control and LOHXD1 enhancer knockdown xenograft. The bar graph show average mitotic nuclei of all the tumors analyzed.

**Materials and method:**

**Cell culture**

Human EwS cell lines RD-ES (obtained from CLS), SK-N-MC, CHLA-10, CADO-ES, prostate cancer cell lines LNCaP, 22RV1, and osteosarcoma U2OS cell lines (obtained from ATCC) were maintained in RPMI-1640 media (Gibco, 11875093) supplemented with 10% FBS (HYC, SH30910.03) and 1% penicillin-streptomycin (Invitrogen, 15140122). All cells were grown at 37°C in a 5% $CO_2$ incubator. For the hypoxia experiment, the cells were incubated in hypoxia chamber of 1% of $O_2$, 5% $CO_2$ at 37°C for the indicated time. All cell lines were tested negative for mycoplasma contamination and authenticated by STR profiling.

**Plasmids**

To generate pcDNA3-LOXHD1-Myc：Full Loxhd1 CDS was cloned from P7 mouse organ of Corti using the following primers: NG-138 and NG-64, then re-amplified using primers containing XhoI and EcorI sites respectively, and cloned into pcDNA3. Myc tag was added by using NEBuilder Hifi 2X mixture with primers: PW-83 and PW-84. (Table S2): To generate NLS-coiled-coil-HA constructs, we amplified the sequence of NLS-Coiled-coil from adult mouse testis cDNA, then ligated to pcDNA3.1(+) vector. NheI and NotI sites were used for the construction. HA tag, NheI and NotI sites were added by PCR primers: NG-191-NheI-CC-S and NG-192-NotI-CC-AS (Table S2). pCDNA-HA-HIF1α was a gift from Dr. Frank S. Lee at UPENN, PA.

**CRISPR-cas9 mediated LOXHD1 enhancer knockout and enhancer silencing**

sgRNAs targeting adjacent to the LOXHD1 GGAA microsatellites were designed with the following website: https://portals.broadinstitute.org/gpp/public/analysis-tools/sgrna-design. For the enhancer region knockout, we inserted two pairs of sgRNAs into lentiCRISPR v2 (Plasmid #52961) backbone. For LOXHD1 repression, two independent sgRNAs were cloned into a lentiviral backbone expressing sgRNA and dCas9-KRAB (Plasmid #71236). Lentivirus was packaged using 2nd generation lentiviral packaging systems using the following protocol. $1x10^6$ HEK-293HT cells were seeded in 10cm plates. Next day lentiCRISPR or lenti-dCas9-KRAB plasmids (4 μg) were co-transfected with pVSVg (1 μg) and pSPAX2 (2 μg) using 7 μl Lipofectamine 2000. Media was collected after 48h and 72h of transfection, centrifuged and passed through 0.45 μm filters to clear of any live cells or debris. Virus were then concentrated by 10X with LentiX concentrator (Takara, 631232) aliquoted and stored at -80°C. RD-ES and SK-N-MC cells were infected at a MOI of 10. We selected for positive cells with puromycin at 1μg/ml for three days to obtain stably knock-out/down cells. Cells within 5 passages after puromycin selection were used for all the experiments.

**shRNA knockdown**

shRNAs against FLI1, and ERG (Penn Core Facilities) were packaged using second-generation lentiviral packaging systems as described above. CHLA-10, RD-ES and SK-N-MC were transduced with shFLI1 lentivirus and CADO-ES cells were transduced with shERG lentivirus. Four days post infection, cells were harvested for qRT-PCR analysis.

**RNA extraction and quantitative RT-PCR**

Total RNA was isolated from cells using the miRNeasy kit (QIAGEN, 217004) and cDNA was synthesized from 1,000 ng total RNA using SuperScript IV First-Strand Synthesis SuperMix (Life Technologies, 18091200). qPCR was performed using Fast SYBRgreen (Life Technologies, 4385612) on a StepOnePlus Real-Time PCR system (Applied Biosystems). Relative expression was calculated using ΔΔCT values normalized against *GAPDH* expression. All primers were designed using primer 3 (http://frodo.wi.mit.edu/primer3/) and synthesized by Integrated DNA Technologies.

**Chromatin immunoprecipitation (ChIP) qPCR and ChIP sequencing**

ChIP was performed using iDeal ChIP-seq Kit for Transcription Factors (Diagenode, C01010170) according to manufacturer's protocol. In brief, SK-N-MC and RD-ES cells of the control and knockdown groups were trypsinized, washed, and crosslinked with 1% formaldehyde in culture medium for 10 min at room temperature. Cross-linking was terminated by the addition of 1/10 volume 1.25 M glycine for 5 min at room temperature followed by cell lysis and sonication (Bioruptor, Diagenode), resulting in an average chromatin fragment size of 200bp. Chromatin equivalent to $5\times10^6$ cells was isolated and incubated with 5 μg antibody overnight at 4 °C (H3-acetyl K27, H3K4me3, H3K9me3, MED1, and IgG (Diagenode). ChIP DNA was isolated by washing and reversal of cross-linking. The eluted DNA was used for SYBRgreen qPCR. The primer sequences used for ChIP-qPCR are provided in Table S2.

10ng of the ChIP DNA was used for ChIP sequencing library preparation following the protocol of TruSeq ChIP library preparation kit (Illumina, IP-202-1012). Briefly, A single “A” nucleotide was added to the 3' ends of the blunt-ended ChIP DNA fragments and then ligated to a unique adapter. The ligation products were purified and selected at the size of 250-300bp by 4% NuSieve agarose gel (Lonza) electrophoresis. Size selected DNA was purified and PCR-amplified and quantitated with the Bioanalyzer 2100 (Agilent). Libraries were pooled and ran on Nextseq500 platform (Illumina Inc.) with high throughput single-end reads of 75bases (Cat.TG-160-2005).

**ChIP-seq analysis**

Sequencing reads for both in-house generated and publicly available datasets were uniformly processed using an in-house ChIP-seq analysis pipeline. Reads were quality checked with FASTQC (www.bioinformatics.babraham.ac.uk/projects/fastqc) and aligned to the GRCh37 (release 27) genome using the STAR v2.5.1 aligner (*1*) with default settings. PCR duplicate reads in the aligned bam files were removed using samtools (*2*), bam files were converted to CPM normalized bigwig tracks using deeptools (*3*), and viewed using the Integrated Genomics Viewer(*4*) or the UCSC genome browser (*5*). Single end reads were extended up to the fragment length (200bp) along the read direction.

**Enrichment analysis of ChIP-seq data**

The enrichment peaks for the various transcription factors and histone marks were computed using MACS2(*6*), with default settings. Regions found to ubiquitously enriched across a number of next-generation sequencing experiments (*7*), also known as the blacklisted peaks (https://sites.google.com/site/anshulkundaje/projects/blacklists), were excluded in all subsequent analysis.

**Overlap of enrichment peaks**

Overlap analysis of enrichment peaks in different samples was performed using an in-house python script. In our analysis, two peaks were taken to be overlapping even if they were a single base.

**RNA-seq library preparation and sequencing**

After the treatments, total RNA was isolated using miRNeasy kit (QIAGEN) and the quality of the RNA was analyzed by Bio-analyzer (Agilent) using RNA nano chip. After confirming that all of the RNA samples have RNA integrity number (RIN) more than 8, RNA-seq libraries were constructed using the TruSeq sample Prep Kit V2 (Illumina) according to the manufacturer's instructions. Briefly, 1 μg of purified RNA was poly-A selected and fragmented with fragmentation enzyme. After first and second strand synthesis from a template of poly-A selected/fragmented RNA, other procedures from end-repair to PCR amplification were performed according to the instructions given in the protocol. Libraries were purified and validated for appropriate size on a 2100 Bioanalyzer DNA 1000 chip (Agilent Technologies, Inc.). The DNA library was quantitated using Qubit and normalized to 4 nM prior to pooling. Libraries were pooled in an equimolar fashion and diluted to a final concentration of 1.8 pM. Library pools were clustered and run on Nextseq500 platform (Illumina Inc.) with single-end reads of 75 bases (Cat.TG-160-2005).

**RNA-seq analysis**

Single-end sequencing reads were demultiplexed using Illumina *bcl2fastq*, quality checked using FASTQC (www.bioinformatics.babraham.ac.uk/projects/fastqc), and aligned to the GRCh37 (release 27) genome using the STAR v2.5.1 aligner,(*1*) with default settings. The read statistics generated by STAR v2.5.1 was used to ensure that the aligned reads in all samples were over 90% of the total reads. The transcripts were assembled using cufflinks and the count and FPKM tables were computed using cuffnorm (*8*). Principal component analysis on the FPKM values was used to cluster the samples for further quality check.

**Differential Gene expression analysis**

Genes differentially expressed between any two sets of treatment conditions were determined using DESeq2 (*9*) (https://bioconductor.org/packages/release/bioc/html/DESeq.html), a statistical tool that employs shrinkage estimates to compute fold changes. In all our calculations, the raw RNA-seq read counts from biological duplicates, for each treatment condition, was used as the input for DESeq2. Heatmaps for differentially expressed genes were generated using in-house python scripts.

**Gene Ontology analysis**

We determined the Gene Ontology (GO) associated with a given set of genes using the python api for g:Profiler (https://biit.cs.ut.ee/gprofiler). The results were plotted using in-house python scripts.

**Gene set enrichment analysis (GSEA)**

Functional class scoring(*10*) of the all differentially expressed genes, against a given gene signature, was performed using the GSEA tool (*11*), developed by the Broad Institute (https://software.broadinstitute.org/cancer/software/gsea/wiki/index.php/Main_Page).

**Immunoblot Analysis**

For immunoblot analyses, cells were lysed in RIPA buffer (Boston Bioproducts, BP-115DG) supplemented with protease inhibitor (Pierce, A32965). Samples for HIF1α immunoblot were harvested in 1x Laemmli sample buffer (Bio-Rad, 1610737). Lysates were boiled in SDS sample buffer (Invitrogen) and 30-50 μg of protein was separated by SDS-PAGE and loaded onto a PVDF membrane. Membranes were blocked for one hour in blocking buffer (Tris-buffered saline, 0.1% Tween (TBS-T), 5% non-fat dry milk) and incubated overnight at 4°C in primary antibody. Blots were washed with TBS-T and incubated with HRP-conjugated secondary antibody for one hour at room temperature. Blots were washed again with TBS-T and visualized after incubation with chemiluminescent substrate (GE Healthcwere). The antibodies used in the study are provided in Table S3.

**Immunoprecipitation**

For immunoprecipitation assay, HEK293T cells were transfected with pCDNA-HA-HIF1α (gift from Dr. Frank S. Lee, UPENN) and pCDNA-Myc-LOXHD1 at a 1 to 2 ratio. 48h post-transfection, cells were incubated in 1% of $O_2$ hypoxia chamber for 6h followed by lysis in IP buffer (20 mM Tris pH7.5, 150 mM NaCl, 1% Triton-X 100, Protease Inhibitor) and sonication. Whole cell lysates (500 μg) were pre-cleaned by incubation with protein G Dynabeads (Life Technologies) for 2h on a rotator at 4°C. 5 μg antibody was added to the pre-cleared lysates and incubated on a rotator at 4°C overnight. Protein G Dynabeads were then added for 2h. Beads were washed four times in IP buffer, containing 300 mM NaCl, and resuspended in 40 μL of 1x Laemmli sample buffer and boiled at 95°C for 5 min for separation of the protein and beads. Samples were then analyzed by SDS-PAGE and western blotting as described above.

**Immunofluorescent staining**

For general staining 100,000 cells were seeded on a coverslip which fits in wells of 24-well plate, the next day, cells were fixed with 4% of paraformaldehyde for 10min at RT. Cells were then washed with PBS and blocked with 10% goat serum and 0.5% Triton 100 in PBS at RT for 60min. Primary antibody of LOXHD1 and/or HA was diluted at 1:500 in 10% of goat serum in PBS and incubated with the cells at 4°C overnight. On the next day, secondary antibody goat anti-Rabbit Alexa Fluor 568 (Thermo Fisher, A-11011) and/or anti-Mouse Alexa Fluor 488 (Thermo Fisher, A28175) was diluted at 1:1000 and incubated at RT for one hour. Coverslips were then mounted and stained for DAPI with anti-fade mounting medium (Vector, H-1200). Images were acquired and processed on a Zeiss confocal microscope (LSM 880).

For F-actin staining, cells were plated in the same way, stained with Phalloidin-iFluor 488 at 1:1000 at room temperature for 1h. Images were acquired and processed on a Zeiss confocal microscope (LSM 880). Cell surface area were measured with ImageJ.

For HA-NLS-coiled coil staining in 293T cells: 293T cells (ATCC, CRL-3216) were plated at a density of $8X10^4$ in per well of a 24-well plate with Ploy-L-Lysine (Sigma-Aldrich, P8920-100ML) coated cover slips the day before transfection. 500ng plasmids were transfected by using FuGENE HD regent. 24h and 48h after transfection, the cells were fixed with 4% PFA for 10 min, RT. The cells were permeabilized and blocked by incubation in blocking buffer (4% BSA, 0.5% saponin in PBS) for 1h at room temperature and then incubated with 1st antibody (rat anti-HA, 3F10, Roche, 1:200 in blocking buffer) overnight at 4°C. The next day, the cover slips were washed with PBS for three times and then incubated in 2nd antibody (Alexa Fluor 488, Goat anti-rat IgG, Thermo Fisher, A11006, 1:300) and DAPI (Sigma-Aldrich, D8417, 30ng/ml) diluted with blocking buffer for one hour at room temperature, followed by 3 times of washing with PBS. Lastly, coverslips were mounted with Prolong Gold anti-fade mounting medium (Invitrogen, P36934). The images were acquired using Zeiss LSM 880 confocal microscope.

**Colony and sphere formation assay**

For colony formation assay, 5,000 cells were plated in one well of the 6-well plates, in two weeks cells were fixed and stained with 0.5% of crystal violet, quantification was done by de-staining of crystal violet and measure absorbance at 560nm. For sphere formation assay, 500 cells were suspended in 100µL 50% of Matrigel of full RPMI culture media and spread on the edge of wells in a 24-well plate, we then fill the wells with 1 ml of full RPMI culture media. In three weeks, spheres were counted, and the sizes of the spheres were measured with ImageJ. Triplicates were carried in all the above described experiments; student t-test was used for statistical analysis.

**Cell aggregation assay**

24-well plates were coated with 100 μL of 3% of poly-HEMA and dried in cell culturing hood overnight. 10,000 cells in single cell suspension were seeded in the wells, pictures were taken at time 0 and 16hr after seeding. Aggregation sizes were measured with ImageJ, student t-test was used for statistical analysis.

**Wound healing assay**

Cells were grown as monolayer in 6-well plate, scratches were made with 1000 μL tips. Pictures of the same area were taken at 24 and 48h. Gap between scratches were measured by ImageJ.

**Matrigel invasion assay**

Stably knockout/down cells were trypsinized and 100,000 cells were suspended in 500μL serum-free RPMI medium and added into Matrigel coated invasion chambers (Corning, 354480). Groups of hypoxic and normoxic samples were performed side by side. The bottom of the chamber was filled with RPMI containing 20% serum as chemo attractant. Cells that had degraded the matrix and migrated through the porous membrane (8 μm pore size) to the other end after a period of 24h or 48h were fixed and stained with crystal violet (0.5%), and images were captured using phase contrast microscopy. The same cells were seeded into chambers with no Matrigel coating as migration control. % invasion was determined by numbers of cells invaded divided by numbers of cells migrated. Invasion index was calculated by % invasion test cells / % invasion control cells. Triplicates were carried in all the above described experiments; we take six images for each well and count the cell numbers. Student t-test was used for statistical analysis.

**Chicken chorioallantoic membrane assay for tumor cell intravasation**

The CAM assay for tumor cell intravasation was performed as previously described (*12*). Briefly, fertilized chicken eggs were incubated in a rotary humidified incubator at 38°C for 10 days. A small hole was drilled through the eggshell into the air sac and another hole was drilled near the allantoic vein that penetrates the eggshell, keeping the chick chorioallantoic membrane (CAM) intact. The CAM was dropped by applying mild vacuum to the hole over the air sac. Subsequently, a cutoff wheel (Dremel) was used to cut a 1 $cm^2$ window encompassing the second hole near the allantoic vein to expose the underlying CAM. Cells were prepared for implantation by trypsinizing and resuspending in media (without FBS) at the density of $2 \times 10^6$ cells/50 μL. The CAM was gently abraded with a sterile cotton swab to provide access to the mesenchyme and 50 μL cell suspension was implanted on top of it. The windows were sealed, and the eggs returned to a stationary incubator. The eggs remained in the incubator for 72h, after which the egg was cut along the long circumference, and the upper half (with the inoculum) and the content of the egg was discarded; the CAM that lines the cavity of the eggshell was lifted and snap-frozen. Genomic DNA from the lower CAM was extracted using PureGene Genomic DNA isolation kit (Gentra-Qiagen) following the manufacturer's protocol and used as a template for human Alu sequence amplification by q-PCR to quantitate the difference in the number of cells between control group and LOXHD1 KD group.

**Zebrafish migration assay**

All procedures on zebrafish (Danio rerio) were approved by the Institutional Animal Care and Use Committee of the University of Pennsylvania. Fertilized zebrafish eggs were incubated at 28 °C in E3 solution and raised using standard methods. Embryos were transferred to E3 solution containing 5 µg/ml protease and 0.2 mM1-phenyl-2-thio-urea (PTU, Sigma) at 24 h post-fertilization to dechorionate the fish embryos and prevent pigmentation, respectively. At 48 h post-fertilization, zebrafish embryos were anesthetized with 0.03% tricaine (Sigma) and transferred to an injection plate made with 1.5% agarose gel for microinjection. Approximately 200–400 mCherry tagged EWS cells suspended in conditioned media supplemented with 0.5 mM EDTA were injected into the perivitelline space of each embryo using a XenoWorks Digital Microinjector (Sutter Instrument). Pre-pulled micropipettes were used for the microinjection (Tip ID 50 µm, base OD 1 mm, Fivephoton Biochemicals). After injection, the fish embryos were immediately transferred to PTU-E3 solution. Injected embryos were kept at 33 °C and examined every day to monitor tumor cell migration using an Olympus Ix81 widefield microscope.

**EwS mouse xenograft**

$3.5\times10^6$ SK-N-MC EwS cells were injected in a 1:1 mix of cells suspended in PBS with Geltrex Basement Membrane Mix (Thermo Fisher) in the right flank of 10–12 weeks old NOD/Scid/gamma (NSG) mice. Tumor diameters were measured every second day with a caliper and tumor volume was calculated by the formula $L\times l^2/2$. Once the first tumor of the control group reached an average volume of 1,500 $mm^3$, all animals of the experiment were sacrificed by cervical dislocation. Other humane endpoints were determined as follows: Ulcerated tumors, loss of 20% body weight, constant curved or crouched body posture, bloody diarrhea or rectal prolapse, abnormal breathing, severe dehydration, visible abdominal distention, obese Body Condition Scores (BCS), apathy, and self-isolation. Animal experiments were approved by local authorities and conducted in accordance with ARRIVE guidelines, recommendations of the European Community (86/609/EEC), and UKCCCR (guidelines for the welfare and use of animals in cancer research).

**Statistical analysis**

Statistical analysis was performed using GraphPad Prism 6 software. For individual comparisons, unpaired Student t test was used and $P < 0.05$ were considered significant. Statistical significance for Kaplan–Meier analysis was determined by the log-rank (Mantel–Cox) test.

**Table S2. List of Oligonucleotide Primers used in this study**

| CRISPR gRNAs | Sequences 5' – 3' |
|---|---|
| GGAA Enhancer KO gRNA_1 FWD | CACCGAGAGAAATTAAAAACAAACA |
| GGAA Enhancer KO gRNA_1 REV | AAACTGTTTGTTTTTAATTTCTCTC |
| GGAA Enhancer KO gRNA_2 FWD | CACCGGAAAAACATCTGCAAGCATC |
| GGAA Enhancer KO gRNA_2 REV | AAACGATGCTTGCAGATGTTTTTCC |
| GGAA dcas9 gRNA_1 FWD | CACCGAGAGAAATTAAAAACAAACA |
| GGAA dcas9 gRNA_1 REV | AAACTGTTTGTTTTTAATTTCTCTC |
| GGAA dcas9 gRNA_2 | CACCGGTAGAGATGACAGGAGTAAA |
| GGAA dcas9 gRNA_2 | AAACTTTACTCCTGTCATCTCTACC |
| **Cloning** | |
| NG-138 | ATGATGGCCCAGAAGAAGAAGCGGAG |
| NG-64 | ACACCCTGCAGCAAGTCCCAACC |
| PW-83 | GAACAAAAACTCATCTCAGAAGAGGATCTTGAGAATTCCACCACACTG |
| PW-84 | TCTGAGATGAGTTTTTGTTCAACGGCCGCGACAGACGGGAAGAGCTC |
| NG-191-NheI-CC-S | CAGCTGGCTAGCACCATGGTGTGGCTGCGGCACCTGGTG |
| NG-192-NotI-CC-AS | TTGCGGCCGCTCAAGCGTAATCTGGAACATCGTATGGGTAAACGGCCGCAATCACCTCCTGCATCCCTGGCC |
| **Genomic PCR validation** | |
| GGAA enhancer FWD | AAGTGGAACTCAGTGTGGAACA |
| GGAA enhancer REV | GCAGGGCACAGAACAGGTACCT |
| **SYBR Green qPCR** | |
| LOXHD1 FWD | TAGTGACCAGGCTGGGACTTG |
| LOXHD1 REV | GCTTCTCCACTTCTATCCCCT |
| FLI-1 FWD | TTAAGGAGGCTCTGTCGGTG |
| FLI-1 REV | GAGGGGGTTGATCTTGTGGG |
| CCK FWD | AGGGTATCGCAGAGAACGGA |
| CCK REV | GGGCCTGCTGGATGTATCTT |
| GAPDH FWD | TGCACCACCAACTGCTTAGC |

| | |
|---|---|
| GAPDH REV | GGCATGGACTGTGGTCATGAG |
| **CHIP qPCR** | |
| GGAA microsatellite FWD | AAACAAATAGCCTGCCCATCAG |
| GGAA microsatellite REV | CCCTCCTTCCTTCCGTGTTT |
| LOXHD1 TSS FWD | CTCAGGTTCCCGCAGGTGT |
| LOXHD1 TSS REV | GGGGCATCATTCTGTCGGC |
| Non-Specific control FWD | ATCCCCCACAACTCCACCTA |
| Non-Specific control REV | ACAGGTAGCAACGAACTGGG |
| **Human Alu Taqman qPCR** | |
| Human Alu FWD | GTCAGGAGATCGAGACCATCCT |
| Human Alu REV | AGTGGCGCAATCTCGGC |
| Human Alu Taqman probe | 5′-6-FAM-AGCTACTCGGGAGGCTGAGGCAGGA-TAMRA-3′ |

**Table S3. List of Antibodies used in this study**

| **Antibody** | **Use** | **Supplier** | **Catalog Number** |
|---|---|---|---|
| H3K27ac | ChIP | Active motif | 39133 |
| H3K4me3 | ChIP | Millipore | 07-473 |
| LOXHD1 | IB/IF | Grillet N, *et al. (2009)* | |
| FLI-1 antibody | IB | Abcam | ab15289 |
| GAPDH-HRP | IB | Cell Signaling | 3683 |
| HIF1α | IB | Cayman Chemical Company | 10006421 |
| HA | IB/IF | Cell Signaling Technology | 3724S |
| Myc-tag | IB/IP | Cell Signaling Technology | 2276S |
| Phalloidin-iFluor 488 Reagent | IF | Abcam | ab176753 |
| Goat anti mouse IgG 594 | IF | Fisher Scientific | A11032 |
| Goat anti rabbit IgG 488 | IF | Fisher Scientific | A11008 |
| Goat anti mouse IgG 488 | IF | Fisher Scientific | A11029 |
| Goat anti rabbit IgG 568 | IF | Fisher Scientific | A11011 |
| Mouse IgG | IP | Diagenode | K01641008 |
| Rabbit IgG | IP | Cell Signaling Technology | 2729S |

**Table S1. Details on ESS32 genes**

| # | ESS32 Gene list | Full name | Gene ontology (biological process) | Function | Status in Ewing sarcoma literature | | |
|---|---|---|---|---|---|---|---|
| 1 | KLF15 | Kruppel Like Factor 15 | positive regulation of transcription by RNA polymerase II [GO:0045944] | Encodes for a transcriptional regulator that binds to, among other promoter regions, the CLCNKA promoter. It is known to inhibit MEF2A and GATA4, thereby playing a role in controlling cardiac hypertrophy. It has also been elucidated as a negative regulator of TP53 acetylation. | Todd M. Stevens, International Journal of Surgical Pathology, 2018 | | |
| 2 | PDE1B | Phosphodiesterase 1B | apoptotic process [GO:0006915] | Cyclic nucleotide phosphodiesterase with a dual-specificity for the second messengers cAMP and cGMP, which are key regulators of many important physiological processes. Has a preference for cGMP as a substrate | None | | |
| 3 | STEAP2 | STEAP2 Metalloreductase | copper ion import [GO:0015677] | Metalloreductase that has the ability to reduce both Fe(3+) to Fe(2+) and Cu(2+) to Cu(1+). | Inês M. Gomes, Molecular Cancer Research, 2012 | Thomas G. P. Grunewald, Biology of the Cell, 2012 | |
| 4 | ABI3 | ABI Family Member 3 | regulation of cell migration [GO:0030334] | The encoded protein is known to inhibit ectopic metastasis of tumor cells and cell migration via interaction with p21-activated kinase. | None | | |
| 5 | DNAJC12 | DnaJ Heat Shock Protein Family (Hsp40) Member C12 | None | Encodes for a member of a subclass of the HSP40/DnaJ protein family, which are known to associate with complex assembly, protein folding, and export. | None | | |
| 6 | PPP1R1A | Protein Phosphatase 1 Regulatory Inhibitor Subunit 1A | intracellular signal transduction [GO:0035556] | Inhibitor of protein-phosphatase 1 | Wen Luo, Nature Oncogene, 2018 | Wen Luo, Nature Oncotarget, 2020 | Daniel H Wai, International Journal of Oncology, 2002 |
| 7 | NPY1R | Neuropeptide Y Receptor Y1 | adenylate cyclase-inhibiting G-protein coupled receptor signaling pathway [GO:0007193] | Receptor for neuropeptide Y and peptide YY. | Jason U. Tilan, Oncotarget, 2013 | | |

| | | | | | | | |
|---|---|---|---|---|---|---|---|
| 8 | VAV1 | Vav Guanine Nucleotide Exchange Factor 1 | regulation of GTPase activity [GO:0043087] | VAV proteins are guanine nucleotide exchange factors (GEFs) for Rho family GTPases, which activate downstream pathways that lead to actin cytoskeletal rearrangements and transcriptional changes. | Rodolphe Guinamard, Scandinavian Journal of Immunology, 1997 | | |
| 9 | DUSP26 | Dual Specificity Phosphatase 26 | protein dephosphorylation [GO:0006470] | Encodes for a tyrosine phosphatase, and has the ability to dephosphorylate both tyrosine and serine/threonine residues. Protein product may regulate neuronal proliferation. This gene has been described as both a tumor suppressor and an oncogene. | None | | |
| 10 | KCNAB3 | Potassium Voltage-Gated Channel Subfamily A Regulatory Beta Subunit 3 | ion transmembrane transport [GO:0034220] | Encodes for protein that forms a heterodimer with the potassium voltage-gated channel, shaker-related subfamily of proteins. | None | | |
| 11 | RBM11 | RNA Binding Motif Protein 11 | cell differentiation [GO:0030154] | Tissue-specific splicing factor with potential implication in the regulation of alternative splicing during neuron and germ cell differentiation. | Andrew J. Annalora, Oncotarget, 2018 | | |
| 12 | KCNE3 | Potassium Voltage-Gated Channel Subfamily E Regulatory Subunit 3 | negative regulation of voltage-gated potassium channel activity [GO:1903817] | Regulates neurotransmitter release, heart rate, insulin secretion, neuronal excitability, epithelial electrolyte transport, smooth muscle contraction, and cell volume. | None | | |
| 13 | MYOM2 | Myomesin 2 | muscle contraction [GO:0006936] | Binds to myosin, titin, and light meromyosin. Shares genetic homology to fibronectin type III and immunoglobulin C2 domains. | None | | |
| 14 | PRR5L | Proline Rich 5 Like | negative regulation of protein phosphorylation [GO:0001933] | Associates with the mTORC2 complex, thereby regulating cellular processes such as survival and cytoskeletal organization. | None | | |
| 15 | NKX2-2 | NK2 Homeobox 2 | positive regulation of transcription by RNA polymerase II [GO:0045944] | Protein-coding gene that contains a homeobox domain and has possible role in the morphogenesis of the central nervous system. | Mitchel J. Machiela, Nature Communications, 2018 | Leah A. Owen, PLoS One, 2008 | Richard Smith, Cancer Cell, 2006 |
| 16 | XG | Xg Glycoprotein | homotypic cell-cell adhesion [GO:0034109] | Encodes for the XG blood group antigen | ynet, Cancer Research, 2010 | | |

| | | | | | | | |
|---|---|---|---|---|---|---|---|
| 17 | RNF182 | Ring Finger Protein 182 | protein ubiquitination [GO:0016567] | Encodes for E3 ubiquitin-protein ligase. Mediates the ubiquitination of ATP6V0C and marks it for degradation through the ubiquitin-proteasome pathway. Inhibits the TLR triggered innate immune response via ubiquitination and subsequent degradation of NF-kappa-B component RELA. | None | | |
| 18 | KCNA2 | Potassium Voltage-Gated Channel Subfamily A Member 2 | potassium ion transport [GO:0006813] | A voltage-gated potassium channel found primarily in the brain, central nervous system, and the cardiovascular system. | None | | |
| 19 | PRRT4 | Proline Rich Transmembrane Protein 4 | None | A protein-coding gene associated with the disease, Zellweger Syndrome. | None | | |
| 20 | NPY5R | Neuropeptide Y Receptor Y5 | cardiac left ventricle morphogenesis [GO:0003214] | Receptor for neuropeptide Y and peptide YY. | Jason U. Tilan, Oncotarget, 2013 | Jason Tilan, Neuropeptides, 2016 | Congyi Lu, The Journal of Biological Chemistry, 2011 |
| 21 | TNNI3 | Troponin I3, Cardiac Type | muscle contraction [GO:0006936] | Part of the Troponin I subfamily of genes. It encodes for the TnI-cardiac protein which is only expressed in cardiac muscle tissues. | None | | |
| 22 | ARTN | Artemin | neuroblast proliferation [GO:0007405] | Encodes for the ligand that activates the GFR-alpha-3-RET receptor complex. | None | | |
| 23 | CD79A | CD7B-Cell Antigen Receptor Complex-Associated Protein Alpha Chain9a | B cell receptor signaling pathway [GO:0050853] | A B lymphocyte antigen receptor that works in conjunction with CD79B to initiate the signal transduction cascade activated by the binding of an antigen to the B-cell antigen receptor complex. | David R. Lucas, Anatomic Pathology, 2001 | Metin Ozdemirli, Nature Modern Pathology, 2001 | Metin Ozdemirli, The American Journal of Surgical Pathology, |
| 24 | PHOSPHO 1 | Phosphoethanolamine/Phosphocholine Phosphatase 1 | bone mineralization involved in bone maturation [GO:0035630] | A phosphatase that has a high specificity for phosphoethanolamine (PEA) and phosphocholine (PCho). Plays a role in generating inorganic phosphate for bone mineralization. | None | | |
| 25 | ADRB3 | Adrenoceptor Beta 3 | adenylate cyclase-modulating G protein-coupled receptor signaling pathway [GO:0007188] | The protein product is part of the beta adrenergic receptor family, and is involved in the regulation of lipolysis and thermogenesis. | Andreas Kirschner, Oncotarget, 2016 | | |

| | | | | | | | |
|---|---|---|---|---|---|---|---|
| 26 | DCDC2 | Doublecortin Domain Containing 2 | cellular defense response [GO:0006968] | Encodes for protein that plays a role in the inhibition of canonical Wnt signaling pathway. | None | | |
| 27 | GNGT2 | G Protein Subunit Gamma Transducin 2 | G protein-coupled receptor signaling pathway [GO:0007186] | Encodes for protein that plays a crucial role in cone phototransduction, and is specifically localized in cones. | None | | |
| 28 | FEZF1 | FEZ Family Zinc Finger 1 | neuron migration [GO:0001764] | Encodes for a transcriptional repressor protein, and is involved in the axonal projection and proper termination of olfactory sensory neurons. | None | | |
| 29 | UGT3A2 | UDP Glycosyltransferase Family 3 Member A2 | cellular response to genistein [GO:0071412] | UDP-glucuronosyltransferases catalyze phase II biotransformation reactions in which lipophilic substrates are conjugated with glucuronic acid to increase water solubility and enhance excretion. | None | | |
| 30 | LOXHD1 | Lipoxygenase Homology Domain-Containing Protein 1 | calcium ion transmembrane transport [GO:0070588] | Involved in hearing. Required for normal function of hair cells in the inner ear. | None | | |
| 31 | LIPI | Lipase I | a potent bioactive lipid mediator) and fatty acid. Does not hydrolyze other phospholipids, like phosphatidylserine (PS), phosphatidylcholine (PC) and phosphatidylethanolamine (PE) or triacylglycerol (TG). {ECO:0000269\|PubMed:12963729}. | Hydrolyzes specifically phosphatidic acid (PA) to produce 2-acyl lysophosphatidic acid (LPA | Juergen L. Foell, Pediatric Blood & Cancer, 2008 | Dorothea E. Mahlendorf, Cancer Biology & Therapy, 2013 | Benjamin J. Schmiedel, Molecular Biology Reports, 2011 |
| 32 | RAX | Retina And Anterior Neural Fold Homeobo | visual perception [GO:0007601] | Contains a homeobox domain, and encodes for a transcription factor known to have functions in eye development. Required for retinal cell fate determination and regulates stem cell proliferation. | None | | |
